# Multisensory integration in *Anopheles* mosquito swarms: The role of visual and acoustic information in mate tracking and collision avoidance

**DOI:** 10.1101/2024.04.18.590128

**Authors:** Saumya Gupta, Antoine Cribellier, Serge B. Poda, Olivier Roux, Florian T. Muijres, Jeffrey A. Riffell

## Abstract

Malaria mosquitoes mate in swarms. Here, they must rely on multiple sensory cues in shaping their individual responses, such as during mate recognition, swarm maintenance, and collision avoidance. While male mosquitoes are known to use faint female flight tones for recognizing their mates, the role of other sensory modalities remains less explored. By combining free-flight and tethered flight simulator experiments with *Anopheles coluzzii*, we demonstrate that swarming mosquitoes integrate visual and acoustic information to track conspecifics and avoid collisions. In tethered experiments, acoustic stimuli gated male steering responses to visual objects simulating nearby female mosquitoes, whereas visual cues alone triggered changes in wingbeat amplitude and frequency. Free-flight experiments show that mosquitoes modulate their flight responses to nearby conspecifics similarly to tethered animals, allowing for collision avoidance within swarms. These findings suggest that combined visual and acoustic information contributes to conspecific recognition within swarms, and, for males, permits female tracking while avoiding collisions.

**Teaser:** Malaria mosquitoes can use sight and sound to navigate swarms, dodging collisions while seeking out mates.

## Introduction

Lekking is a remarkable behavioral strategy documented across various species, in which individuals congregate at specific areas – termed leks – to engage in competitive displays with the ultimate aim of securing a mate (*1*, *2*). This strategy, widely studied in birds and mammals, is also paralleled in the insect world, where swarms formed for the purpose of mating function as analogous structures to leks (*3*). However, unlike the static gatherings seen in many lekking species, flying insects such as dance flies, mayflies, midges and mosquitoes form highly dynamic mating swarms (*4*–*6*). In these swarms, individuals navigate through a complex three-dimensional space, introducing the challenges of aerial maneuvering to an already complicated in-flight mating ritual. Successful mating within these swarms thus hinges upon the ability of individuals to adeptly avoid collisions in the complex sensory environment while pursuing mates (*7*). *Anopheles* mosquitoes, known for their role in the transmission of malaria-causing pathogens, are one group of swarm-forming insects whose mating success holds significant consequences for public health (*8*, *9*). Yet, the sensorimotor mechanisms that regulate the interplay of avoidance and attraction among conspecifics within these swarms remain largely unknown.

The swarming behavior of the *Anopheles gambiae s.I.* mosquitoes, representing the best characterized mating behavior among mosquitoes (*10*–*12*), serves as an excellent model for understanding the sensorimotor mechanisms operating within swarms. As dusk falls, mosquitoes of this species complex begin to aggregate over visual landmarks, forming swarms that serve as mating arenas. These swarms are predominantly composed of males, who compete for the relatively few females in the swarms. Males actively pursue females for copulation while managing to avoid collisions with other males, presumably by maintaining a specific inter-individual distance (*7*). Such spatial navigation within swarms suggests an advanced sensory mechanism at play, one that is critical yet not fully understood. Several studies have revealed that males rely on their acute ability to detect the distinct flight tones of females, which are produced at a different frequency than their own, to identify and pursue potential mates (*13*–*16*). On the contrary, males are much less sensitive to the flight tones of other males and may not always be able to acoustically detect one another (*17*–*21*). This selective acoustic sensitivity suggests the potential existence of other sensory mechanisms beyond auditory processing that facilitate individual detection and collision avoidance within swarms.

Previous research have established that mosquitoes can integrate multiple sensory inputs, including visual, thermal, and olfactory cues (*22*–*25*). For example, host-associated olfactory cues can trigger mosquitoes to initiate visual search behaviors, even in nocturnal ones like *Anopheles* (*24*, *26*). This capability to simultaneously process and integrate multiple sensory cues is believed to be crucial for host-seeking activities and is likely instrumental in other behavioral contexts as well. An example of the potential integration of multiple sensory modalities, in another behavioral context than host-seeking, comes from swarming mosquitoes that use visual markers to form and maintain swarms and acoustic signals from female wingbeats to locate mates within these swarms (*27*–*29*). However, the extent to which swarming behavior is guided by the integration of these visual and acoustic cues remains an open question. Within swarms, where hundreds to thousands of buzzing mosquitoes occupy a fixed volume, the resulting sensory landscape rich in acoustic and visual information suggests that mosquitoes may also use their visual system for detecting fellow swarming individuals (*30*). Whether mosquitoes employ their visual system along with auditory system to interact with conspecifics within the dynamic setting of swarms remains to be tested.

In this study, we tested the hypothesis that *Anopheles* mosquitoes utilize visual cues of flying individuals in conjunction with acoustic information from their wingbeats to approach potential mates and avoid collisions with conspecifics. This hypothesis challenges the widely held belief that *Anopheles* mosquitoes rely minimally on visual cues for detecting other individuals due to their poor visual acuity, estimated at an angular resolution of 16.7° (*31*). However, it is important to recognize that flying insects, despite their low spatial resolution, are generally well-adapted to visually detect moving targets representing prey, predator or mates (*32*). Supporting this, a study on *Aedes aegypti* has demonstrated mosquitoes’ capability to detect small moving objects (*33*). Given the close proximity of individuals within swarm (*34*, *35*), it stands to reason that *Anopheles* mosquitoes are also likely capable of detecting the motion of other individuals, particularly those within short range.

We investigated our visual-acoustic integration hypothesis using a combined tethered-flight and free-flight experimental approach. In our tethered experiments, we used a virtual reality flight simulator to examine the behavioral response of rigidly tethered *Anopheles coluzzii* to moving objects, in the presence or absence of conspecific sounds. In our free-flight experiments, we examined free-flight responses of *Anopheles coluzzii* in swarms and compared these with responses in the tethered state. The rigidly tethered mosquito in a flight simulator allowed for fine-scale control of the statistics of the visuo-acoustic environment – not permitted in other tethered preparations or during free-flight. While such simulators cannot replicate the complexity of natural swarming conditions, they are invaluable for dissecting the fundamental aspects of insect sensory behavior under tightly controlled settings (*36*–*38*). Our work, leveraging these approaches, reveals the importance of sensory integration in shaping conspecific interactions within mosquito mating swarms.

## Results

### *Anopheles* mosquitoes respond to swarm-like visual scenes

Our initial experiment focused on assessing whether *Anopheles* mosquitoes could potentially detect aspects of visual environment characteristic of swarms. Given the limited angular resolution of mosquito eyes, detecting the fine details of swarms are likely beyond their visual capacity (*31*). However, the high light sensitivity of their eyes might enable them to detect the contrast of aggregated mosquitoes against the twilight sky (*31*). To explore this idea, we simulated the visual environment of mosquito aggregations on a two-dimensional cylindrical LED panel, displaying random starfield patterns with 10% of the LEDs illuminated. This design reflected the density of contrasting elements visible against the backdrop of the sky, as observed in the natural swarm imagery (Fig. 1A).

**Fig. 1:**
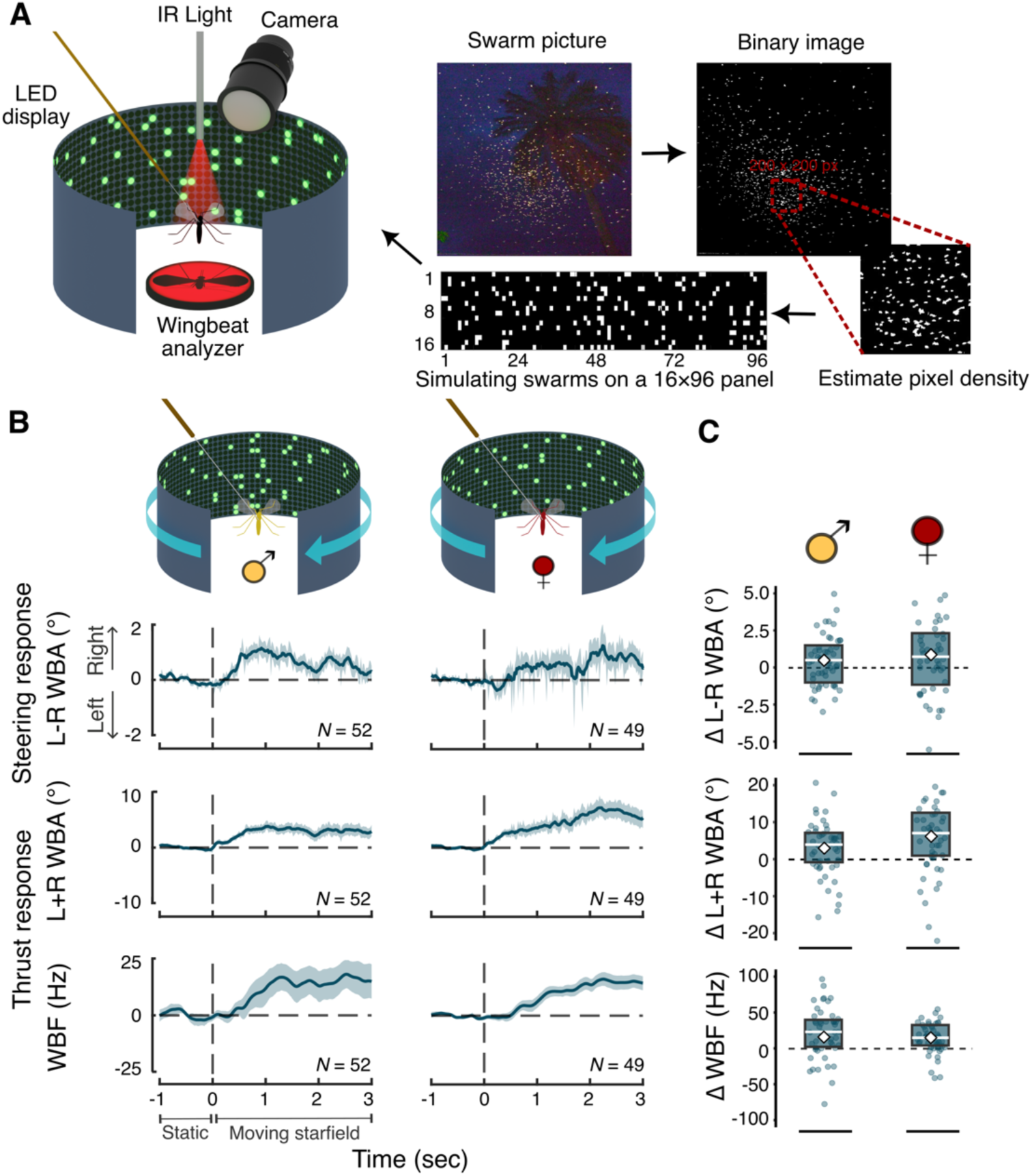
*Anopheles* mosquitoes behaviorally respond to simulated swarm-like visual scenes. (**A**) Illustration of the visual flight simulator, which includes a LED panel for presenting visual stimuli and a camera and wingbeat analyzer to record wing kinematics of tethered mosquitoes. The panel displays starfield pattern derived from binarized images of actual *Anopheles* swarms. Each pixel subtends an angle of 3.75° on the mosquito eye. The pattern aims to replicate the contrast density of mosquitoes observed in natural swarms against the background of the sky. (**B**) Mean normalized behavioral responses of male (left) and female (right) *An. coluzzii* to visual stimuli. Shaded regions represent the standard error (± SE). Top panel shows steering responses measured by difference in left and right wingbeat amplitude, while the middle and bottom panels display additional thrust response measured by total wingbeat amplitude and wingbeat frequency, respectively. The data is averaged across both clockwise and counterclockwise motion and standardized for clockwise rotation (**C**) Statistical analysis of male (left) and female (right) responses, shown via boxplots, quantifies the change in response from the one second of pre-stimulus baseline (static) to the last one second of the stimulus presentation (moving starfield). Boxplot elements include the interquartile range (box boundaries), mean (diamond), median (horizontal line), with individual data points that fall within at least 95% quantile range. Mean and median symbols are color coded with white symbols denoting statistical significance (*P* < 0.05). In this experiment, all measured responses during stimulus presentation significantly differed from baseline.

To evaluate whether mosquitoes can detect these simulated swarm-like scenes, we presented both male and female mosquitoes with static starfield patterns and monitored their behavioral response as the pattern drifted horizontally, in clockwise or counterclockwise (yaw) directions. Typically, a moving visual environment induce steering behaviors in tethered flying insects, characterized by differential wingbeat amplitudes of the left and the right wings (L-R WBA) indicative of steering, as well as increase in the total wingbeat amplitude (L+R WBA) and wingbeat frequency (WBF), which together are defined as contributing to an increase in wingbeat-induced aerodynamic thrust (*33*, *39*–*41*).

Our analyses revealed that both male and female mosquitoes responded to the visual stimuli by steering towards the direction of the moving starfield patterns (Fig. 1B *top row*). Compared to their baseline response when the starfield pattern was static, both sexes exhibited a significant steering towards the direction in which the pattern was moving (Fig. 1C *top row*; males: *P* = 0.034, *t* = 2.18, *df* = 51; females: *P* = 0.025, *t* = 2.31, *df* = 48, Paired t-test). Responses were similar for both clockwise and counterclockwise motion after accounting for the direction (males: *P* = 0.73, *ß* = 0.17, SE = 0.50, *df* = 26; females: *P* = 0.14, *ß* = -0.96, SE = 0.63, *df* = 25, Linear mixed effect model), supporting averaging the responses to both stimulus motion directions for comprehensive analysis. This behavior was coupled with changes in aerodynamic thrust production, as evidenced by increased total wingbeat amplitude (Figs. 1B &C *middle row*; males: *P* = 0.004, *t* = 3.02*, df* = 51; females: *P* < 0.001, *t* = 4.15, *df* = 48) and wingbeat frequency (Figs. 1B &C bottom row; males: *P* = 0.017, t = 2.47, *df* = 51; females: *P* < 0.001, t = 5.09, *df* = 48). Our comparative analysis showed no significant differences in steering and thrust responses between males and females (L-R WBA: *P* = 0.404, *t* = -0.84, *df* = 79.72; L+R WBA: *P* = 0.083, *t* = -1.76, *df* = 85.03; WBF: *P* = 0.901, *t* = 0.12, *df* = 71.43, Welch Two Sample t-test). These results collectively highlight the potential of *Anopheles* to be able to detect aspects of the visual environment of natural swarms.

### Acoustic cues modulate visual object tracking

Next, we investigated whether visual cues could play a role in shaping conspecific interactions, particularly in the contexts of mate attraction and collision avoidance. Based on our hypothesis that *Anopheles* mosquitoes combine visual and acoustic information, we predicted that mosquitoes would exhibit acoustically dependent visual response. Without a conspecific flight tone present, mosquitoes would show aversion from visual objects, indicative of collision avoidance. In contrast, if the same object is accompanied by a flight tone of a potential mate, the mosquitoes would be attracted towards the visual objects. Given mosquitoes’ antennae and sensory neurons are not tuned to detecting the flight tones of same sex individuals (*13*, *42*, *43*), we also expected to observe aversive behaviors from visual objects paired with same-sex flight tones. To test these predictions, we focused on evaluating the steering response of male and female mosquitoes since it is a reliable method of measuring attraction or repulsion in tethered flying dipterans (*44*–*46*).

Our setup involved simulating the visual cue of a nearby flying mosquito within a swarm on our LED panel. We displayed a square object with a 22.5° optical angle, moving against a dark starfield background (Fig. 2A). This optical angle approximates the angular size of a mosquito viewed by another mosquito from a distance of 2.4 body lengths. To further add the biological context of whether the visual object represents a male or a female mosquito, we designed additional stimulus presentations in which the moving object was concurrently broadcast with species-specific flight tones (female-like flight tone at 450 Hz and male-like flight tone at 700 Hz) (Fig. 2A).

**Fig. 2.**
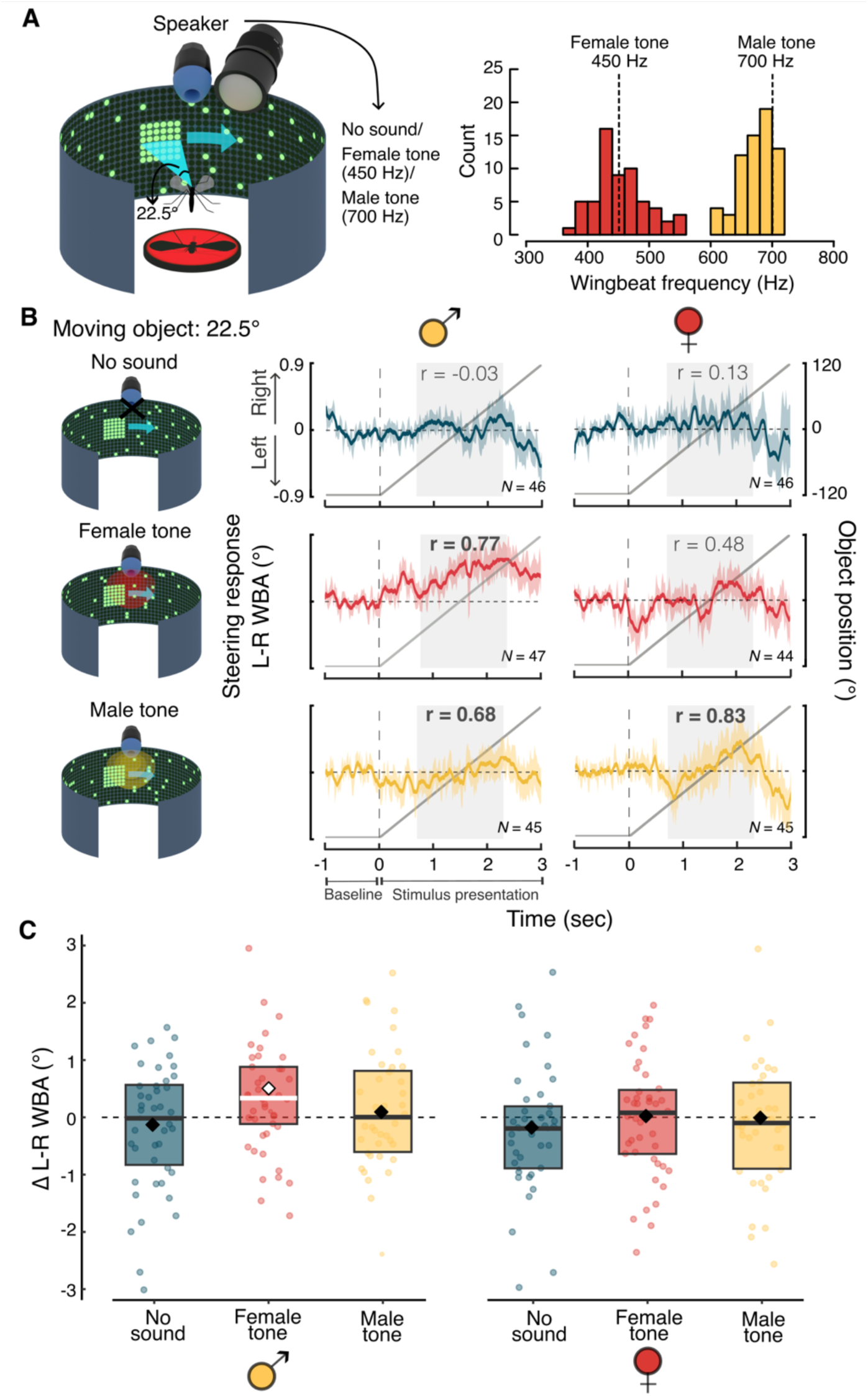
Acoustic modulation of object tracking in males and females. (**A**) Illustration of the tethered flight setup used in this experiment. A square object against a starfield pattern, subtending at angle of 22.5° at the mosquito’s eye, simulates a visual cue of a nearby mosquito within a swarm. An earphone, fixed in front of the tethered mosquito, delivers either a female-like tone at 450 Hz or a male-like tone at 700 Hz [these values are based on previous literature (*39*, *40*) and the distribution of tethered female (red) and male (yellow) wingbeat frequencies recorded when mosquitoes were not presented with any visual or acoustic cue]. (**B**) Mean normalized steering responses of tethered male (left) and female (right) *An. coluzzii* to the 22.5° square object during a pre-stimulus baseline condition when the object was static (-1 to 0 sec) and during the stimulus presentation when object moved horizontally (0-3 sec), with or without accompanying acoustic cues. Responses are averaged for over both directions of object motion and standardized to represent motion from left to right. Shaded regions represent standard error (± SE). The solid gray line indicates the location of object, with 0° representing it directly in front of the mosquito. r value on each plot indicates Spearman’s correlation between average steering response and the object position during the time segment highlighted by light gray shaded region. Values highlighted in bold signify strong correlation (r > 0.6). The full dynamics of steering responses before, during, and after stimulus presentation can be seen in Fig.s S2 and S3. (**C)** Statistical analysis of male (left) and female (right) responses, shown via boxplots, quantifies the change in response from a pre-stimulus baseline to stimulus presentation. Boxplot elements include the interquartile range (box boundaries), mean (diamond), median (horizontal line), with individual data points that fall within at least 95% quantile range. Mean and median symbols are color coded with white symbols denoting statistical significance (*P* < 0.05).

In supplemental experiments, we showed that similar to free-flying males (*29*, *47*), tethered males exhibited steering responses towards the direction of female flight tone (Supplementary Fig. S1). Therefore, in these set of experiments we fixed the position of the speaker in front of the tethered mosquito to ensure that all responses were due to visual stimuli. This experimental design also resulted in our decision to use a rigidly tethered setup over a magnotether system (*48*, *49*), which would have allowed for free rotation of mosquitoes around the yaw axis but would have altered the relative orientation of the head to the sound source.

During each stimulus presentation, we monitored the steering responses of individuals as the object moved horizontally (yaw) from one side to the other. Monitoring the steering responses allowed us to assess two aspects of object tracking behavior in mosquitoes. Firstly, we examined changes in steering response from a pre-stimulus baseline when the object was static to the period when the object was moving, which helped evaluate whether mosquitoes were attracted to, or repelled by, the object. Secondly, we analyzed the correlation between the object’s changing position and the mosquitoes’ steering responses, which provided insights into their spatial awareness regarding the object.

Contrary to our expectation, and different from responses to the star-field (wide-field) motion, mosquitoes did not exhibit aversion or attraction towards the moving object when presented alone. Compared to their baseline response, neither males nor females steered away or even towards the moving object in the absence of acoustic cues (Fig. 2B *top row* & Fig. 2C; males: *P* = 0.611, *t* = -0.51, *df* = 45; females: *P* = 0.382, *t* = 0.89, *df* = 45, Paired t-test) and their steering responses were also not correlated with the location of the object (males: r = -0.03; females: r = 0.13; Spearman’s correlation). However, in the presence of female tones, males exhibited marked attraction to the object, evident from their significant steering in the direction of the object’s motion (Fig. 2B *middle left column* & Fig. 2C; *P* = 0.012, *t* = 2.60, *df* = 46). In contrast, the presence of male tones did not elicit significant steering towards the moving object (Fig. 2B *bottom left column* & Fig. 2C; *P* = 0.739, *t* = 0.33, *df* = 44), though a strong correlation was observed between the males’ steering behavior and the object’s location during male tone presentation (r = 0.68; Fig. 2B *bottom left column*), which is indicative of visual object tracking, albeit without a directional approach towards it.

Unlike males, females did not alter their steering response in the presence of female flight tones. Compared to their baseline response, they did not exhibit significant steering towards or away from the moving object paired with a female tone (Fig. 2B *middle right column* & Fig. 2C; *P* = 0.972, *t* = -0.03, *df* = 43), and their steering response was also not strongly correlated with the position of the object (r = 0.48). In the presence of male tones, although females did not exhibit significant steering towards the object (Fig. 2B *bottom right column* & Fig. 2C; *P* = 0.896, *t* = -0.13, *df* = 44), their steering responses were very strongly correlated with the location of the moving object (r = 0.83; also refer to Fig. S2), indicating that females did track visual objects in the presence of male tones.

These results reveal that *Anopheles* mosquitoes integrate visual and acoustic information and highlight a potentially important role of visual cues in shaping behavioral dynamics during conspecific interactions.

### Acoustic modulation of visual object tracking depends on object size

Building on our findings that acoustic cues can trigger *Anopheles* mosquitoes, especially males, to track a visual object, we further investigated whether these responses varied with object size. Given that the perceived size of a mosquito changes with distance, understanding this relationship is crucial in the natural context. Therefore, we presented male mosquitoes with square objects of 37.5°, 22.5°, and 15°, which represent the size of mosquitoes at distance of 1.3, 2.4 and 3.7 body lengths, respectively. These angular sizes were selected to align with the visual resolution capabilities of this species and fall within a range that is relevant for inter-individual interactions within swarms.

We observed that the responses to 37.5° object were similar to responses to the previously tested 22.5° object: in the absence of acoustic cues, males did not significantly steer in response to the visual cue (Fig. 3A *top column1&2*; Table 1). However, with the addition of female flight tones, they exhibited a strong steering response in the direction of the object’s motion (Fig. 3A *middle column1&2*; Fig. 3B; Table 1). The male flight tones did not elicit a similar steering behavior in males as female tones (Fig. 3B; Table S1), although a strong tendency to track the object’s location was still observed (r = 0.80, Fig. 3A *bottom column1&2*; also refer to Fig. S3).

**Fig. 3.**
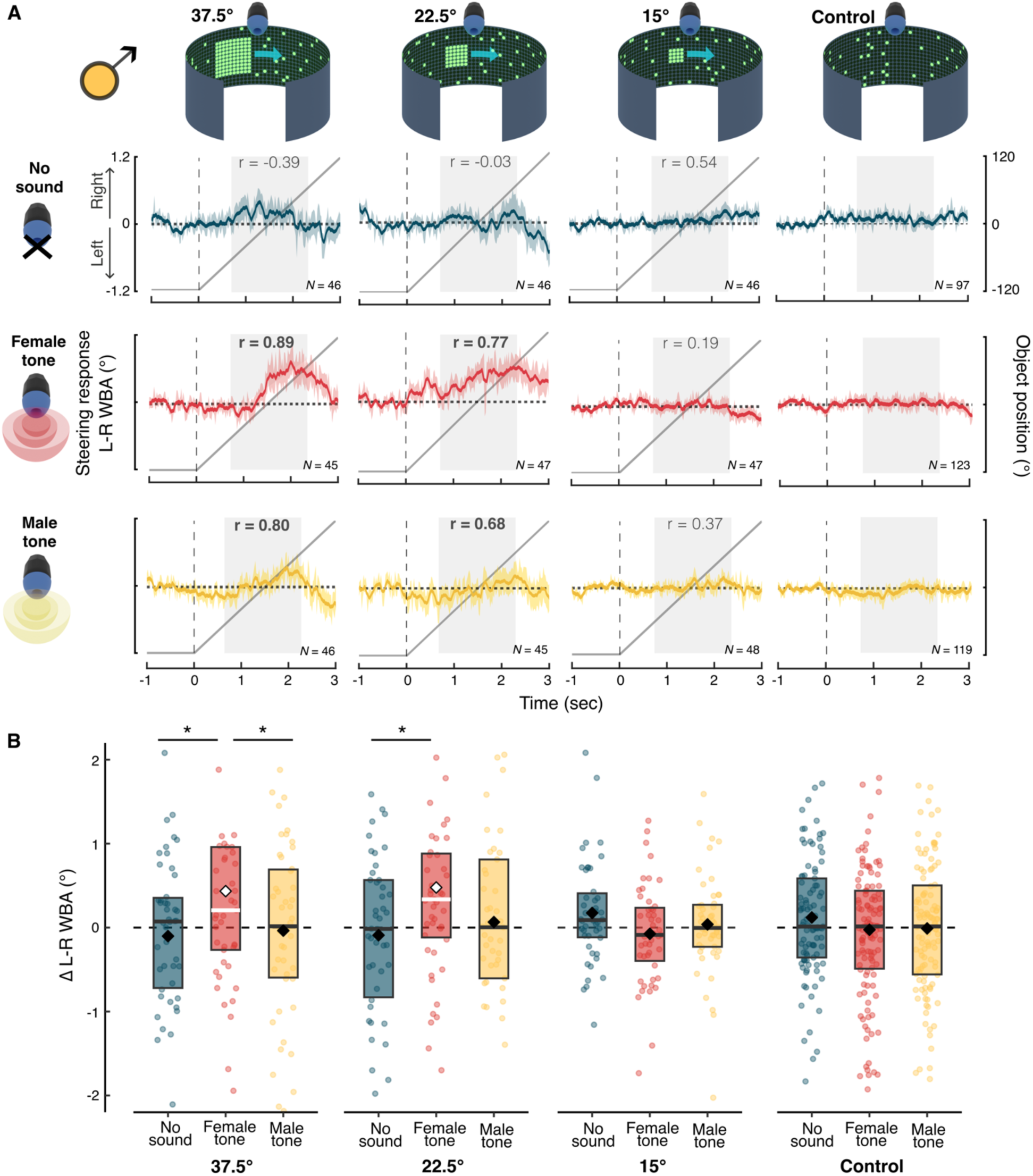
Size dependent acoustic modulation of visual tracking in males. (**A**) Mean normalized steering responses of males to horizontally moving objects of three different sizes (37.5°, 22.5° referenced from Fig. 2 for comparison, and 15°). For each object size, responses are measured under three conditions: without acoustic cue (row 1), in the presence of a female tone (row 2), and in the presence of a male tone (row 3). The solid gray line indicates the position of object, with 0° representing it directly in front of the mosquito. r value on each plot indicates Spearman’s correlation between average steering response and the object position during the time segment highlighted by light gray shaded region. Values highlighted in bold signify strong correlation (r > 0.6). A control condition (column 4) with only the starfield pattern (no moving object) with and without acoustic cues were also tested. The full dynamics of steering responses before, during, and after stimulus presentation for each stimulus treatment can be seen in supplementary Fig. S3. (**B**) Statistical analysis of responses, shown via boxplots, quantifies the change in steering response from a pre-stimulus baseline to stimulus presentation. Boxplot elements include the interquartile range (box boundaries), mean (diamond), median (horizontal line), with individual data points that fall within at least 95% quantile range. Mean and median symbols are color coded with white symbols denoting statistical significance. Asterisks (*) denote significant differences (*P* < 0.05) between treatments when *P-values* are adjusted for multiple comparisons.

In contrast, males showed no steering response to the smaller 15° object, even when it was coupled with acoustic cues (Fig. 3A *column3*; Table 1). These responses were indistinguishable from the control conditions where a static starfield, without any moving object, was presented with and without acoustic cues (Fig. 3A *column4*; Fig. 3B; Table S2). Together, these results reveal a size-dependent threshold in mosquito visual-acoustic integration, suggesting that acoustic stimuli gate target tracking of visual objects, such as other mosquitoes, only when they are in very close-proximity.

**Table 1:** Results of paired *t-tests* comparing steering responses (L-R WBA) of males before and during stimulus presentation.

| **MALES** |  |  |  |  |  |  |  |  |
| --- | --- | --- | --- | --- | --- | --- | --- | --- |
| **37.5° object in static starfield background** |  |  |  |  |  |  |  |  |
| No sound |  |  | Female tone* |  |  | Male tone |  |  |
| *t* = -0.69 | *df* = 45 | *P* = 0.490 | *t* = 2.27 | *df* = 44 | *P* = 0.028 | *t* = -0.24 | *df* = 45 | *P* = 0.811 |
| **22.5° object in static starfield background** |  |  |  |  |  |  |  |  |
| No sound |  |  | Female tone* |  |  | Male tone |  |  |
| *t* = -0.51 | *df* = 45 | *P* = 0.611 | *t* = 2.61 | *df* = 46 | *P* = 0.012 | *t* = 0.33 | *df* = 44 | *P* = 0.739 |
| **15° object in static starfield background** |  |  |  |  |  |  |  |  |
| No sound |  |  | Female tone |  |  | Male tone |  |  |
| *t* = 1.92 | *df* = 45 | *P* = 0.062 | *t* = -0.84 | *df* = 46 | *P* = 0.405 | *t* = 0.43 | *df* = 47 | *P* = 0.666 |
| **Control: Static starfield background** |  |  |  |  |  |  |  |  |
| No sound |  |  | Female tone |  |  | Male tone |  |  |
| *t* = 1.58 | *df* = 96 | *P* = 0.118 | *t* = -0.28 | *df* = 122 | *P* = 0.777 | *t* = -0.11 | *df* = 118 | *P* = 0.911 |

Asterisks (\*) denote significant differences ( $P < 0.05$ )

### Mosquitoes adjust their wing kinematics in response to visual cues

Contrary to our initial expectation that mosquitoes would steer away from visual objects as a collision avoidance response, our observations did not support this behavior. This led us to further examine the effect of visual cues on mosquito responses beyond steering. We expanded our analysis to encompass changes in wing kinematics, specifically total wingbeat amplitude (L+R WBA) and wingbeat frequency (WBF), as males were presented with visual objects of varying sizes in the presence and absence of acoustic cues. Wingbeat amplitude and frequency modulations allow flying insects to adjust the aerodynamic thrust forces required for performing flight maneuvers (*50*–*52*), and thus this investigation aimed to determine whether visual information could influence other flight behaviors that can potentially contribute to collision avoidance strategies.

Our analysis revealed a distinct pattern of modulation in wing kinematics. Males exhibited a decrease in both wingbeat amplitude and frequency as the visual object approached their frontal field of view, indicated by a negative correlation between the object’s location and aerodynamic thrust production (Fig. 4A *column 1-3*). Conversely, a monotonic increase in wingbeat amplitude and frequency was observed as the object moved away, indicated by a strong positive correlation in the subsequent temporal window (Fig. 4A *column 1-3*). This response pattern was observed regardless of whether the visual objects were presented with or without acoustic cues (cf. Fig. 4A *top, middle, and bottom rows*) and was notably distinct from the control condition (static starfield without any object), where the responses remained unchanged or decreased monotonically (Fig. 4A *column 4*).

**Fig. 4.**
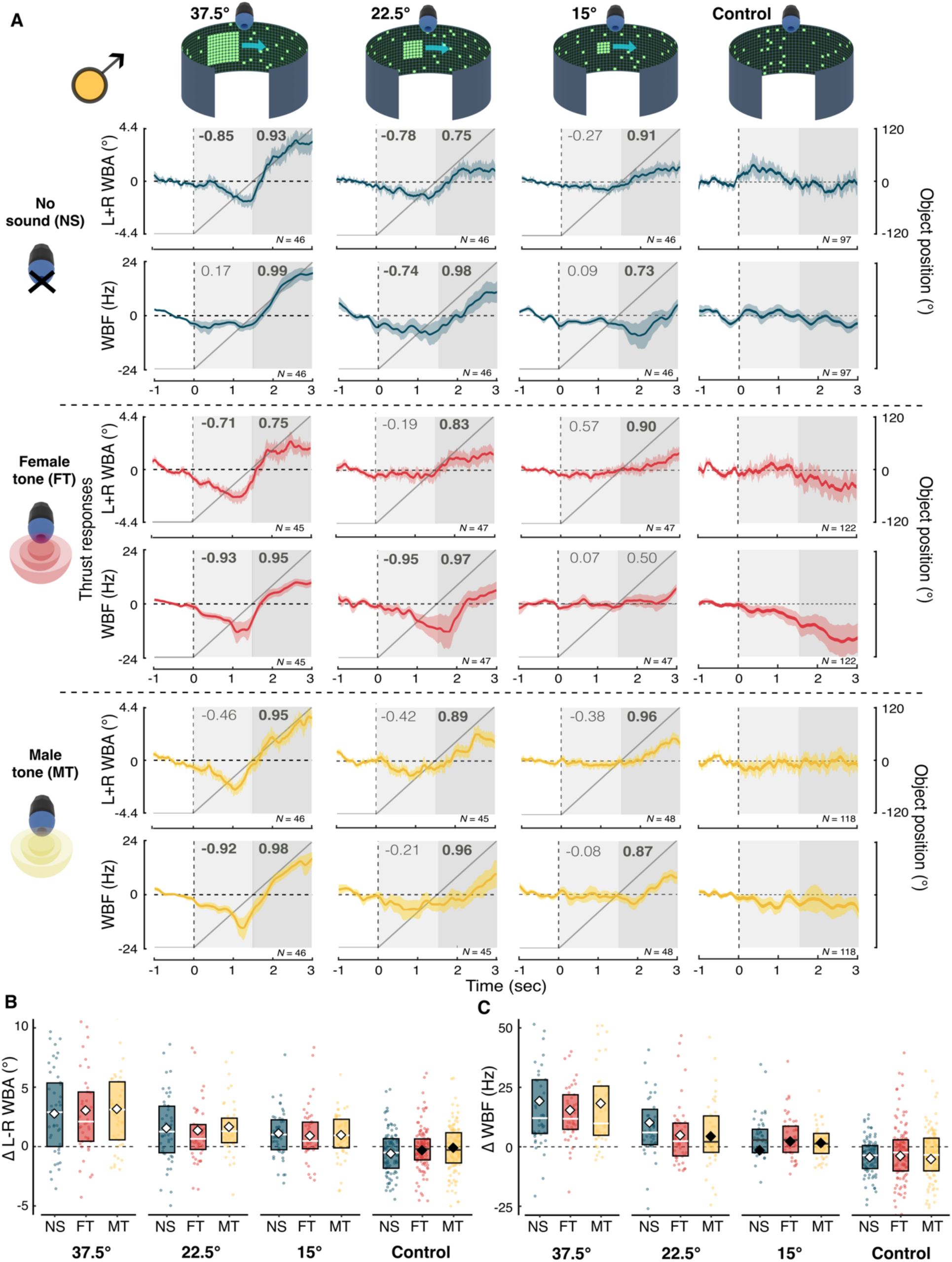
Modulation of male wingbeat kinematics in response to object motion. (**A**) Normalized mean thrust responses of male mosquitoes, as indicated by total wingbeat amplitude (L+R WBA) and wingbeat frequency (WBF) in response to visual objects of three different sizes (37.5°, 22.5°, 15°). For each object size, responses are measured under three conditions: without acoustic cue, in the presence of a female tone, and in the presence of a male tone. The solid gray line indicates the position of object, with 0° representing it directly in front of the mosquito. r value on each plot indicates Spearman’s correlation between average steering response and the object position during the time segment highlighted by light gray shaded region. Numbers at the top of the gray colored segments indicates Spearman’s correlation between average wing kinematics and the object position during the time segments highlighted by gray shaded regions. Values highlighted in bold signify strong correlation (r > 0.6). A control condition (column 4) with only the starfield pattern (no moving object) with and without acoustic cues were also tested. The full dynamics of wing kinematics before, during, and after stimulus presentation for each object size across the acoustic treatments can be seen in supplementary Fig.s S4 and S5. (**B**) Statistical analysis of wingbeat amplitude responses, and (**C**) wingbeat frequency responses shown via boxplots, quantifies the difference in response across acoustic treatments as the object moves to the front (0 to 1.5 sec) and as it moves away from the front (1.5 to 3 sec). Boxplot elements include the interquartile range (box boundaries), mean (diamond), median (horizontal line), with individual data points that fall within at least 95% quantile range. Mean and median symbols are color coded with white symbols denoting statistical significance.

The observed modulation in wingbeat amplitudes in relation to the object’s position was statistically significant across all acoustic treatments (no sound, female tone, male tone) and object sizes (37.5°, 22.5°, and 15°), confirming that this was a visually driven behavior (Fig. 4B; Tables 2 & S3; Fig. S4). The modulation in wingbeat frequency was also similar to wingbeat amplitude modulations across all object sizes and treatments, though significant changes in wingbeat frequency were limited to objects of 37.5° and 22.5° (Fig. 4C; Table 3; Fig. S5). While the changes in wingbeat frequency for the 15° object were not statistically significant, the modulation pattern was still significantly different from the no-object control condition (Tables S4). Notably, the extent of modulation in thrust production, resulting from the combined wingbeat amplitude and frequency adjustments, varied with object size, with the most pronounced changes occurring in response to the largest (37.5°) visual stimulus (Fig. 4B&C, Tables S3 & S4). "

**Table 2:** Results of *t-tests* evaluating whether changes in male wingbeat amplitude (L+R WBA) during stimulus presentation are significant.

| **37.5° object in static starfield background** |  |  |  |  |  |  |  |  |
| --- | --- | --- | --- | --- | --- | --- | --- | --- |
| *No sound\** |  |  | *Female tone\** |  |  | *Male tone\** |  |  |
| *t* = 4.87 | *df* = 45 | *P* < 0.001 | *t* = 4.53 | *df* = 44 | *P* < 0.001 | *t* = 5.31 | *df* = 45 | *P* < 0.001 |
| **22.5° object in static starfield background** |  |  |  |  |  |  |  |  |
| *No sound\** |  |  | *Female tone\** |  |  | *Male tone\** |  |  |
| *t* = 3.49 | *df* = 45 | *P* = 0.001 | *t* = 3.07 | *df* = 46 | *P* = 0.004 | *t* = 4.86 | *df* = 44 | *P* < 0.001 |
| **15° object in static starfield background** |  |  |  |  |  |  |  |  |
| *No sound\** |  |  | *Female tone\** |  |  | *Male tone\** |  |  |
| *t* = 3.61 | *df* = 45 | *P* < 0.001 | *t* = 2.27 | *df* = 46 | *P* = 0.028 | *t* = 3.42 | *df* = 47 | *P* = 0.001 |
| **Control: Static starfield background** |  |  |  |  |  |  |  |  |
| *No sound\** |  |  | Female tone |  |  | Male tone |  |  |
| *t* = -2.87 | *df* = 96 | *P* = 0.005 | *t* = -1.37 | *df* = 122 | *P* = 0.173 | *t* = -0.68 | *df* = 118 | *P* = 0.497 |

Asterisks (\*) denote significant differences ( $P < 0.05$ )

**Table 3:** Results of *t-tests* evaluating whether changes in male wingbeat frequency (WBF) during stimulus presentation are significant.

| **37.5° object in static starfield background** |  |  |  |  |  |  |  |  |
| --- | --- | --- | --- | --- | --- | --- | --- | --- |
| *No sound\** |  |  | *Female tone\** |  |  | *Male tone\** |  |  |
| *t* = 5.78 | *df* = 45 | *P* < 0.001 | *t* = 6.65 | *df* = 44 | *P* < 0.001 | *t* = 6.11 | *df* = 45 | *P* < 0.001 |
| **22.5° object in static starfield background** |  |  |  |  |  |  |  |  |
| *No sound\** |  |  | *Female tone\** |  |  | Male tone |  |  |
| *t* = 2.95 | *df* = 45 | *P* = 0.005 | *t* = 2.26 | *df* = 46 | *P* = 0.029 | *t* = 1.36 | *df* = 44 | *P* = 0.181 |
| **15° object in static starfield background** |  |  |  |  |  |  |  |  |
| No sound |  |  | Female tone |  |  | Male tone |  |  |
| *t* = -0.47 | *df* = 45 | *P* = 0.643 | *t* = 0.78 | *df* = 46 | *P* = 0.437 | *t* = 0.538 | *df* = 47 | *P* = 0.593 |
| **Control: Static starfield background** |  |  |  |  |  |  |  |  |
| *No sound\** |  |  | *Female tone\** |  |  | *Male tone\** |  |  |
| *t* = -3.83 | *df* = 96 | *P* < 0.001 | *t* = -2.35 | *df* = 122 | *P* = 0.020 | *t* = -2.30 | *df* = 118 | *P* = 0.023 |

Asterisks (\*) denote significant differences ( $P < 0.05$ )

Subsequent experiments on female *An. coluzzii*, involving only visual stimuli without acoustic cues, revealed a similar modulation pattern across all three object sizes (Fig. S6). Together, these results suggest that mosquitoes are capable of maneuvering in response to visual objects by adjusting their wingbeat kinematics, potentially utilizing this sensorimotor mechanism as a means to avoid collisions.

### Collision avoidance and tracking behavior of male mosquitoes flying freely in swarms

Our controlled virtual arena experiments suggest that tethered mosquitoes modify their flight patterns in response to visual objects simulating other mosquitoes at close range. Specifically, they change their wingbeat amplitude and frequency in response to visual objects and modulate their steering behavior based on whether the visual object is paired with flight tones from a male or a female conspecific. To understand if these behavioral patterns found in tethered experiments also reflect free-flight patterns in natural swarming scenarios, we analyzed free-flight dynamics within *An. coluzzii* swarms (Fig. 5A) (*53*). Given that the swarms of this species are predominantly male and it is challenging to identify the sex of individual mosquitoes in a swarm using current videography methods, our laboratory swarms were composed entirely of males. Across the six swarming events recorded in (*53*), the mosquito densities were low, with only 13 to 27 males swarming over a 40×40 cm ground marker. The average distance to the nearest neighbor in this dataset was 12.23 cm, with instances of close encounters within 2 cm constituting less than 1% of all nearest-neighbor distances within the swarm (Fig. 5B). Considering that the wingspan of *Anopheles* mosquitoes is roughly 0.6 cm (*54*), another mosquito within 2 cm would subtend a visual angle greater than 16.7°. This particular measure of proximity is especially relevant as it provides a benchmark to correlate the flight patterns observed in the controlled tethered setting with behaviors manifested during free flight.

**Fig. 5.**
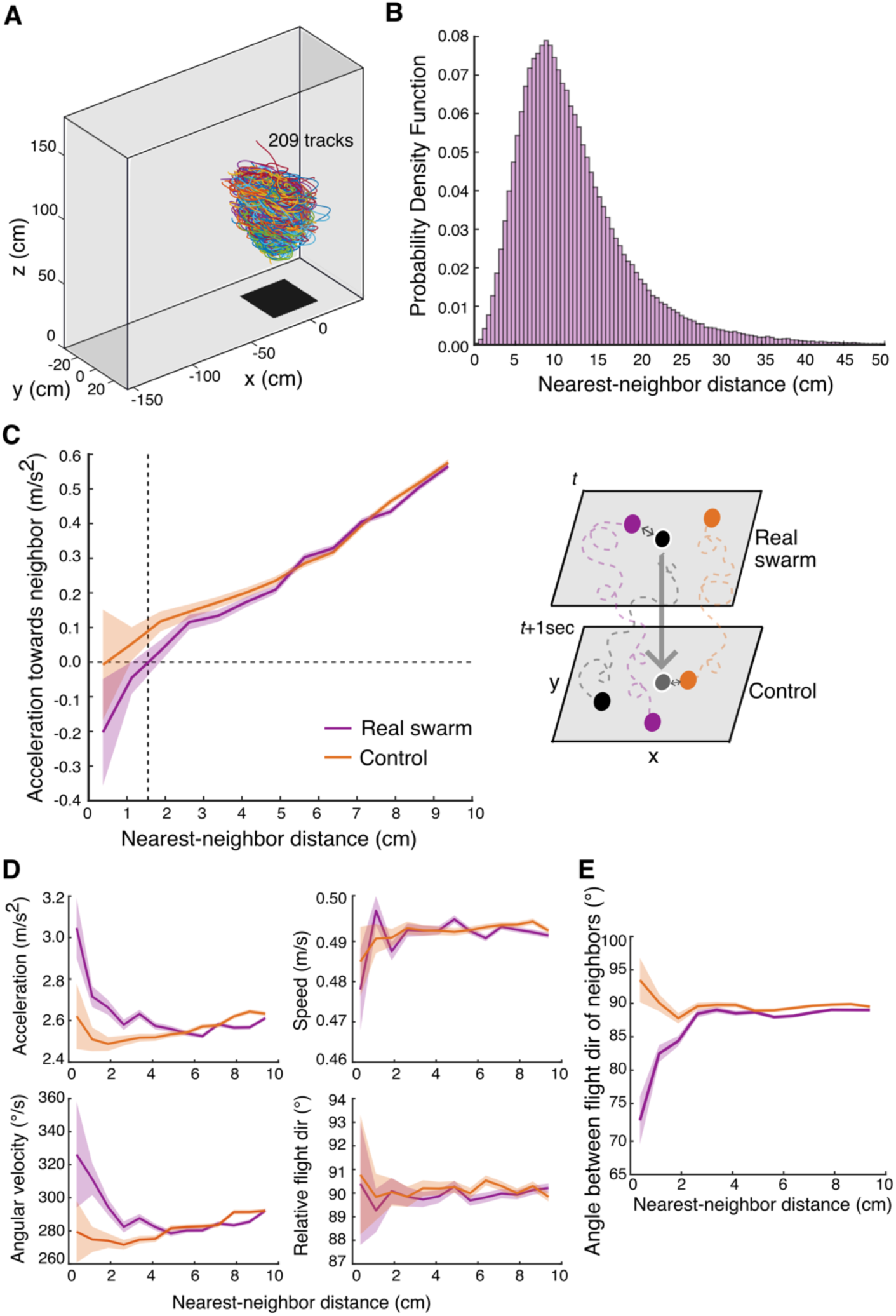
Analysis of flight dynamics as a function of nearest-neighbor distances in male swarms. (**A**) Flight trajectories of male *An. coluzzii* within a laboratory-generated swarm. The illustrated swarming event includes 25 mosquitoes, recorded for 150 seconds and resulting in 209 recorded tracks. (**B**) Probability density function of nearest-neighbor distances across the six swarms (938 total trajectories). (**C**) *Left*. Plot of acceleration of mosquitoes towards their nearest neighbor as a function of distance. Acceleration here is calculated by projecting the focal mosquito’s acceleration onto the direction of its nearest neighbor, measuring the extent of repulsive and attractive forces between mosquitoes. Real swarm data (with *N* = 311631 frames in which nearest-neighbor pairs were calculated are represented in purple, while control data (with *N* = 304657 frames) are in orange. *Right*. Schematic representation of real swarm interactions (focal mosquito in black at time *t* with its nearest neighbor in purple) and hypothetical interactions in control data (focal mosquito projected to time *t*+1sec in gray with its new nearest neighbor in orange at *t*+1sec). (**D**) Relationship between various flight parameters of mosquitoes (acceleration, angular velocity, flight speed, and flight direction relative to its nearest neighbor) as a function of nearest neighbor distances. (**E**) Angle between flight direction of neighboring mosquitoes, measured as the angle between their velocity vectors, across different nearest-neighbor distances.

First, we looked for evidence of collision avoidance in our dataset. Based on our tethered experiments, we expect mosquitoes may prevent collisions with visual objects by adjusting their wingbeat amplitude and frequency, resulting in a modulation of the aerodynamic thrust force. We therefore sought evidence for change in such forces when freely swarming mosquitoes flew in close-proximity to each other. Inspired by research on other swarming species like midges and schooling fish in which interactions between individuals in a social group were modelled as effective forces (*55*, *56*), we looked at whether mosquitoes tend to attract or repel each other by measuring the instantaneous acceleration of each mosquito towards its nearest neighbor. Here, positive and a negative acceleration would signify attraction towards and repulsion from the nearest neighbor, respectively.

Our analysis indicated a clear ‘repulsion zone’: on average, when mosquitoes were within 1.55 cm of each other (∼2.5 body lengths), they accelerate away from their neighbor (negative accelerations), suggesting an intrinsic mechanism to prevent collisions (Fig. 5C). Conversely, at larger distances, we noted a pattern of increasing acceleration towards the nearest neighbor, which is a characteristic pattern that emerges when swarming individuals remain in the swarm (*56*). To validate our findings, we performed a hypothetical experiment using the same swarming dataset, but whereby we projected each focal mosquito one second into the future (Fig. 5C). This allowed us to create a control dataset consisting of flight dynamics of a virtual mosquito in relation to its apparent nearest neighbors. Analysis of this control dataset revealed a positive average acceleration of virtual mosquitoes towards their apparent neighbors at all distances, and thus on average no repulsion was observed from the control (Fig. 5C). Notably, at neighbor distances smaller than 4 cm, the mosquitoes in the real data showed significantly lower accelerations towards their neighbors than the control (0-2 cm: *P* = 0.007, *ß* = -0.11, *SE* = 0.04; 2-4 cm: *P* = 0.036, *ß* = -0.03, *SE* = 0.12, *df* = 615346, Linear mixed effect model). In contrast, at distances larger than 4 cm, accelerations were not significantly different between the real and control mosquitoes (4-6 cm: *P* = 0.303, *ß* = -0.011, *SE* = 0.01, *df* = 615346). Together, this analysis shows that the mosquitoes actively avoid collisions during close encounters, but did not respond to conspecifics at distances larger than 4 cm.

Continuing our examination of free-flight dynamics, we specifically investigated how mosquitoes might be evading collisions. Our analysis indicated that during close encounters, mosquitoes increased both their acceleration and angular velocity, suggesting that they performed rapid turning maneuvers (Fig. 5D *left column*). Statistical comparison with the control scenario validated that the observed increase in acceleration and angular velocity was a direct result of interaction with neighbors in close proximity (acceleration: 0-2 cm: *P* < 0.001, 2-4 cm: *P* < 0.001, 4-6 cm: *P* = 0.339, *df* = 615346; angular velocity: 0-2 cm: *P* < 0.001, 2-4 cm: *P* < 0.001, 4-6 cm: *P* = 0.636, *df* = 613071; refer to Table S5 for other model parameters). Interesting, flight speed during close encounter remained consistent across nearby mosquitoes in the real and control scenarios (Fig. 5D *right column*; 0-2 cm: *P* = 0.247, 2-4 cm: *P* =0.358, 4-6 cm: *P* = 0.791, *df* = 615346; Table S5), suggesting that mosquitoes adjusted their flight path rather than speed to evade each other.

Mosquitoes and fruit flies have been shown to evade looming objects using directional evasive maneuvers (*57*–*59*). However, our results also suggest that sharp directional changes away from the neighbor were not a factor in the avoidance strategies in the swarm; real mosquitoes did not significantly alter their flight direction relative to the position of their nearest neighbor compared with virtual mosquitoes (Fig. 5D *right column*; 0-2 cm: *P* = 0.431, 2-4 cm: *P* = 0.331, 4-6 cm: *P* = 0.186, *df* = 615346; Table S5). These patterns appear to be in line with our observations from tethered flight experiments suggesting that swarming mosquitoes might prevent collisions not through sharp directional changes, but rather through adjustments in aerodynamic thrust forces, potentially enhancing maneuverability.

In our final analysis, we sought evidence in free flight that corroborates our tethered experiment finding that male mosquitoes’ steering responses are also influenced by the visual-acoustic representation of a nearby male. Supporting this, we found that the neighboring mosquitoes adjusted their flight directions when within 2 cm (Fig. 5E). Beyond this proximity, the average angle between flight directions of neighboring mosquitoes converged to the average angle of approximately 90°, suggesting that neighboring mosquitoes at larger distances flew in random directions (Fig. 5E; (0-2)-(2-4) cm: *P* < 0.001, *ß* = -5.52, *SE* = 0.80; (2-4)-(4-6) cm: *P* = 0.127, *ß* = 0.650, *SE* = 0.35, *df* = 350950). In the virtual control scenario, this adjustment in flight direction at close neighbor distances was absent (0-2 cm: *P* < 0.001, *ß* = -5.89, *SE* = 1.06, *df* = 350950; Table S5), as neighboring flight directions were largely unrelated regardless of nearest neighbor distances (Fig. 5E; (0-2)-(2-4) cm: *P* = 1.000, *ß* = -0.54, SE = 0.81; (2-4)-(4-6) cm: *P* = 0.166, *ß* = 0.79, *SE* = 0.36, *df* = 350950). These results parallel our observations from tethered experiments, hinting at a possible vigilance mechanism that may enable males to keep track of nearby males without direct interaction.

## Discussion

For swarming mosquitoes, acoustic sensing has long been the focal point of our understanding of inter-individual interactions, possibly overshadowing the importance of other sensory cues. This study broadens the current perspective by exploring the potential role of vision in shaping conspecific interactions within mating swarms. We discovered that *An. coluzzii* mosquitoes can detect and respond to visual objects representative of other mosquitoes, but their visual response is modulated by the sex-specific flight tones. Notably, males exhibited a pronounced directional response towards visual objects associated with female tones, indicating the integration of visual and acoustic signals to locate females. Additionally, we found that visual cues alone were sufficient to trigger changes in wingbeat kinematics, potentially aiding in collision avoidance through subtle flight modifications. Many of the behavioral patterns we found in tethered experiments were supported by free-flight dynamics of swarming mosquitoes, providing compelling evidence in support of our hypothesis that *Anopheles* mosquitoes utilize visual cues of flying individuals in conjunction with acoustic information to approach potential mates and avoid collisions with conspecifics.

A growing body of research has documented the importance of visual cues for mosquito in various contexts, including host localization (*23*, *25*, *26*), swarm formation (*27*, *28*) and threat avoidance (*58*, *60*). However, no published study has investigated the potential contribution of visual cues to mating behaviors and interactions with other conspecifics. Our work has filled this gap by demonstrating that swarm-forming mosquitoes can not only respond to visual cues of conspecifics within mating swarms but also integrate these cues with acoustic signals to facilitate mating interactions. The observed behavioral contrast in our results – where males exhibited no notable steering towards visual objects in the absence of sound but a marked steering towards objects in the presence of female sounds – indicates that acoustic information critically gates the visual response.

Sex-specific differences is another dimension to consider in the acoustic gating of visual behavior of mosquitoes. Although, our data on females are comparatively limited, the differential response between sexes are noteworthy. In contrast to males, females did not exhibit tracking behavior towards a visual object paired with a female tone. They displayed object tracking only in response to male tones; however, their response appeared different from the male’s directed response towards objects representing potential mates. These observations align with the known differences in auditory sensitivity (*14*, *61*), which suggest that although both sexes can detect flight tones of potential mates, males possess specialized mechanism that amplifies female flight tones, enhancing their sensitivity to these particular sounds. These findings add complexity to the interplay of acoustic and visual cues and reinforce the integral role of visual tracking in mosquito mating behaviors.

We interpret our findings on visuo-acoustic integration to suggest that mosquitoes use visual cues of other flying mosquitoes as a supplementary aid alongside acoustic cues for interacting with nearby individuals, especially potential mates. Specifically, we propose that male mosquitoes, after locating a potential mate by her flight tones, are likely to hone in on her precise location using visual cues as she becomes visually discernible. This sensory strategy is particularly relevant when males rapidly modulate their wingbeat frequency while chasing females (*18*, *62*) – a behavior that can reduce the detectability of female flight tones due to changes in distortion product generation (*21*). Distortion products, arising due to the nonlinear mixing of male and female flight tones within the mosquito’s auditory system, are known to mediate acoustic detection of females (*19*, *63*, *64*). Thus, as a male mosquito changes its wingbeat frequency, its ability to detect these distortion products, and thereby the female, also changes, potentially making visual cues crucial for accurate localization. Furthermore, females may also utilize visual cues to orient themselves towards a nearby male, without necessarily approaching the male, reflecting the well-established mating dynamics where males actively pursue and intercept females (*65*, *66*). Finally, our findings suggest that males may also rely on combined visual and acoustic information to track other males at close distances, supporting the notion of male-male interactions beyond simple avoidance, as previously suggested (*67*–*69*).

The distinct integration of visual and acoustic stimuli in *An. coluzzii* not only highlights a sophisticated behavioral adaptation but also raises intriguing questions about the underlying neural mechanisms. Given that the auditory system of male mosquitoes is more sensitive to detecting female flight tones compared to male tones (*13*, *19*, *70*), and that the degree of modulation in steering behavior observed in our experiments corresponded with the acoustic sensitivities, it is reasonable to assume that information processing by the auditory system significantly influences mosquitoes’ visual responses. The precise manner in which acoustic inputs modulate visual processing pathways presents an intriguing subject for future research. Existing literature suggest that neuromodulators are critical to shaping the output of insect neural circuits and altering their behavioral states (*71*–*73*). In mosquitoes, the neuromodulator octopamine has been shown to play an integral role in auditory function (*74*), providing efferent feedback to the Johnston’s organ via octopaminergic neurons (*52*). Interestingly, these neurons also project to the lobula region of the optic lobe, where ‘object selective’ neurons known to process small-field visual information are typically present in insects (*33*, *62*, *63*). This convergence of auditory and visual neural pathways, and innervation by neuromodulator-expressing cells, suggests a potential neural basis for visual-auditory integration (*30*). Investigating the neural pathways and the role of neuromodulators across sensory systems could provide insights into the mechanism of sensory integration, potentially offering novel avenues for vector control strategies.

Beyond the integration of visual and acoustic cues, our study also sheds light on the independent influence of small visual objects on mosquito behavior. Distinct from responses in other Diptera – such as rigidly tethered *Drosophila melanogaster*, which steer away from objects smaller than 30° (*44*, *77*), and *Aedes aegypti*, which steer towards these objects (*33*) – *An. coluzzii* demonstrates a unique response. Instead of differentially modulating their left and right wingbeat amplitude to steer in yaw direction, *Anopheles* mosquitoes modulated other wingbeat kinematics, specifically total wingbeat amplitude and frequency, in response to the visual objects. This modulation was not dependent on acoustic stimuli and varies with the size of the visual object, suggesting that this is a purely visual driven mechanism that mosquitoes might be using to avoid collisions with nearby mosquitoes in a swarm. High-speed videography of escape maneuvers of these mosquitoes reveals that they can rapidly accelerate away from looming threats by increasing wingbeat kinematics to increase acceleration and by simultaneously rotating their body along the pitch and roll axes for turning (*59*). Results from our free-flight analyses revealing that mosquitoes exhibit higher acceleration and angular velocities when close to other mosquitoes, without significant changes in flight direction in relation to the nearest neighbor, hints that they might avoid collisions by rapidly increasing acceleration and making subtle directional shifts facilitated by changes in body rotations. Although we could not measure body rotations due to our tethered setup, the wing kinematic changes in response to visual stimuli may mirror the first phase of this escape maneuver, suggesting that mosquitoes may use similar mechanisms to evade both large, looming objects and small visual objects, such as nearby mosquitoes. Future research aimed at quantification of mosquito kinematics in response to visual object motion during free flight using high-speed videography will be crucial to confirm the proposed role of visual cues in collision avoidance strategies.

In conclusion, this study broadens our understanding of mosquito sensory ecology by highlighting the potential roles of visual cues and interplay between visual and acoustic cues in shaping their interactive behaviors. Our findings offer compelling evidence that mosquitoes actively process and integrate visual and acoustic information, a discovery that has implications for both ecological research and public health initiatives, particularly in the context of vector control. Notably, previous interventions targeting male mosquito swarms have been linked to reduced female insemination rates and decreased malaria prevalence, highlighting the potential of swarm-based interventions in vector management (*8*, *78*). Yet, the practical application of this strategy has been often limited by reliance on insecticides or labor-intensive manual trapping methods. Acoustic lures have emerged as promising alternative for mass male capture, but their effectiveness under natural conditions has shown inconsistent results (*79*–*81*). Our insights into the integration of sensory cues suggest that augmenting acoustic lures with attractive visual stimulus could potentially enhance their effectiveness, providing a more efficient method for capturing swarming mosquitoes.

## Materials and Methods

### Subjects

We used 4- to 6-day-old unmated, wild-type *Anopheles coluzzii* (Ngousso strain, MRA-1279) as test subjects for the experiments with tethered mosquitoes. The colonies were reared in environment control chambers (Intellus, Pervical Scientific Inc., Perry, IA, USA), maintained at a temperature of 26±1°C and 60±10% relative humidity under a 12-hour light/12-hour dark cycle. To ensure that subjects remain unmated, we isolated individual pupae in 50-mL Falcon tubes each day. Upon emergence, adults were provided with cotton saturated with a 10% sucrose solution. Given that this species typically initiates swarming around dusk, all experiments were conducted with during the first hour of the dark cycle to align with their natural swarming patterns. Furthermore, our controlled sensory experiments specifically included mosquitoes that displayed consistent flying while being tethered, ensuring their physiological state closely resembled that during swarming.

### Tethering procedure

Subjects were anesthetized by brief exposure to cold for no more than 3 minutes before being placed on a Peltier cooling stage, maintained at approximately 4°C, to ensure they remained immobilized during tethering. Each individual was subsequently fixed by the thorax to a 0.025-cms wide tungsten rod using UV-activated glue (Loctite 3104 Light Cure Adhesive, Loctite, Düsseldorf, Germany). Prior to testing, the tethered mosquitoes were rested upside down in a transparent container lined with moist paper towels to avoid dehydration, and allowed to recover for at least 20 minutes.

### Virtual reality flight simulator with acoustic setup

The flight responses of tethered mosquitoes to acoustic and visual stimuli were measured in a flight simulator arena, situated in a dark room maintained at 25±1°C. The simulator’s core was a cylindrical LED array featuring a 96×16-pixel grid, with each pixel subtending 3.75° on the mosquito’s eye at the azimuth. The flight simulator has been used in *Drosophila* and now mosquito research (*33*, *45*, *82*–*84*). Subjects were secured in a fixed position within the arena at a 45° pitch angle and positioned under an infrared diode. The position of the mosquito was adjusted until the shadow of the beating wings was detectable by an optical wingbeat analyzer situated underneath the subject (Fig. 1A). This analyzer converted the shadow’s movements into quantifiable data, capturing both the wingbeat amplitude (WBA) and wingbeat frequency (WBF) for analysis.

Challenges arose in measurements from wingbeat analyzer due to the shallow stroke angle of mosquito wings (∼ 40°), yielding extremely noisy data when computing differences in left and right WBA (*58*). This contrasts with the larger differences in left and right WBA observed during steering in other species, such as Drosophila, which exhibit stroke angles near 140° (*85*). To enhance data accuracy, we supplemented our setup with a Firefly USB 2.0 camera (60 frames/s; Model FMVU-03MTM-CS, Point Grey Research Inc., Richmond, BC, Canada) with an infrared (IR) filter (Midwest Optical System, Inc., Palatine, IL, USA) and employed Kinefly, an open source machine-vision software utilizing an edge detection algorithm to detect wingbeat amplitude (*86*). This system enabled more reliable tracking of mosquitoes’ WBA in each video frame. However, we observed that some mosquitoes exhibited excessive leg movements, which interfered with the edge detection of wing strokes. Such individuals were excluded from subsequent analysis.

For the delivery of close-range acoustic stimuli, we integrated the audio output from a computer (Dell Optiplex 7080, Dell Technologies, Austin, TX, USA) equipped with a sound card (Audiophile 2496, M-Audio, Cumberland, RI, USA) into the setup. The audio was amplified using a headphone amplifier (DAC-X6, FX-Audio, Shenzhen, China) and delivered to the tethered mosquitoes via one channel of a stereophone (KOSS Corporation, Milwaukee, WI, USA) positioned directly in front of the subject above the LED panels (Fig. 2A). Acoustic cues were presented from the front because sounds traveling along the horizontal plane are effective at being detected by mosquitoes than sounds traveling in the vertical plane (*87*). To verify that the sounds were playing during audio stimulus delivery, the unused stereophone channel was placed outside the simulation arena along with a calibrated particle velocity microphone (Knowles NR-23158, Ithaca, NY, USA) coupled to an audio interface (M2 interface, MOTU, Cambridge, MA, USA) for digitizing sound input. All data, including the sound stimulus from the stereophone, wingbeat amplitude from the camera, and wingbeat frequency from the wingbeat analyzer, were recorded at a sampling rate of 2000 Hz, which exceeds the Nyquist frequency for the male wingbeat frequency. Data acquisition was facilitated by a National Instrument Acquisition board (BNC 2090A, National Instruments, Austin, Texas, USA) and captured using WinEDR v4.0.0 software (University of Strathclyde, Glasgow, UK). The entire flight simulator setup was enclosed in a custom-made sound isolation box (26.5×26.5×28 cm) to prevent sound reflection and interference from external noise. The box was also fitted with a thick acoustic curtain, which was drawn closed during experimental runs for additional sound insulation.

### Simulating swarm-like visual scenes on LED panels in the flight simulator

To simulate the visual scenes characteristic *An. coluzzii* swarms, we analyzed fifteen photographs taken approximately 2 meters from the center of natural swarms. The locations where the images were taken have been sites of previous research on mosquito swarming behaviors in Burkina Faso, West Africa (*8*, *88*, *89*). These images were processed into binary format, highlighting the mosquitoes as white spots against a black background. We selected a uniform area of 200×200 pixels within regions of high swarm density to determine the percentage of white pixels corresponding to area occupied by mosquitoes. The observed pixel densities varied from 1.6% to 9.1%, averaging at 4.4%. To simulate the visual scenes of swarms on our LED panel, we chose a pixel density of 10%, aligning with the higher end of the natural swarm densities to ensure biological relevance and adequate visual contrast.

Our simulation involved the generation of random starfield patterns on a 96×16 pixel LED panel, using bright pixels set against a dark background to represent the visual contrast in a swarm. This visual environment was also effective at imitating the light intensity characteristic of twilight conditions (0.003 W/m^2^ in our setup versus 1-0.0006 W/m^2^ from sunset to full lunar illumination (*90*). During experimental trials, we presented subjects (both males and females) with wide-field motion of this starfield pattern in open-loop for three seconds, either in clockwise or counter-clockwise direction, and evaluated their tethered flight responses. In a set of preliminary trials, we also tested the response of mosquitoes to wide-field motion of dark pixels on a bright background; however this stimuli failed to elicit a response, suggesting that such visual conditions provided insufficient contrast detection by the mosquitoes.

### Moving visual objects with species-specific acoustic cues

To simulate visual aspect of close encounters between mosquitoes within a swarm, we generated square objects in three sizes corresponding to the apparent size of a conspecific at varying distances: 37.5° (10×10 pixels), 22.5° (6×6 pixels), and 15° (4×4 pixels). These sizes were selected to align with or exceed the known visual resolution limits of these mosquitoes (*31*) and approximate the sight of a neighbor at 1.3, 2.4, and 3.7 body lengths, respectively. Set against a background of the previously described starfield pattern, the brightly pixelated objects were designed to be visually detectable against the contrasting dark environment.

To complement these visual cues with the acoustic cue of nearby flying conspecific, we generated pure tones mimicking the fundamental wingbeat frequency of a female and a male *An. coluzzii*. The sounds were synthetically generated in MATLAB and presented at a particle velocity of about 0.04 mm/s, which is about the sound intensity generated by this species at a distance of 2 cm (*62*). Such pure tones have been found to be effective at eliciting natural, acoustically guided behaviors in mosquitoes (*87*, *91*).

Different group of subjects were assigned to be tested with one of the three object sizes. Multiple open-loop trials were conducted to assess their response to the moving visual object both in isolation and coupled with the male or female flight tones. Prior to the trial’s onset, a stationary object within the static starfield background was displayed for two seconds. The trial started with the object’s horizontal movement from the left to the right or right to the left visual field, completing two rotations in 6 sec. At the trial’s conclusion, the object was held static for an additional two seconds, allowing subjects to return to their baseline flying patterns. The objects moved at an angular velocity of approximately 80°/sec. During acoustic playback trials, the movement of the visual stimulus was synchronized with a six-second flight tone that matched either the male or female flight tones (700 Hz and 450 Hz, respectively). To ensure that any observed changes in tethered flight behavior were due to moving visual objects, most subjects were also tested in control trials with and without acoustic cues where the visual stimulus was solely the static starfield background. Each subject was presented with these varied trials four times in a pseudorandom order to mitigate any potential order effects. The initiation of each trial type was uniquely coded to ensure precise identification during analysis.

### Tethered flight data processing

Raw data was analyzed using a custom script in MATLAB R 2023a. First, data were down sampled to 50 Hz to align with the camera’s limited frame rate. We then computed additional metrics of differential wingbeat amplitude (L-R WBA) and total wingbeat amplitude (L+R WBA) at each frame using the recorded left and right wingbeat amplitudes. For each trial, baseline measurements of L-R WBA, L+R WBA and wingbeat frequency (WBF) were determined by averaging the values over a one-sec window preceding the start of the trial. Normalized responses for each trial were calculated by subtracting this baseline from the responses recorded during the entire trial period.

Trials where mosquitoes ceased to fly – evidenced by a wingbeat frequency drop below 350 Hz – or exhibited spurious data, as indicated by weak correlation in wingbeat patterns between the left and the right wings (< 0.6), were excluded from analysis. To standardize the data, responses to stimuli moving from right to left were adjusted to reflect responses to left to right motion. The mean normalized response across time was then calculated for each treatment (rotating starfield or moving visual object, moving visual object+female sound, moving visual object+male sound) and control group (static starfield, static starfield+female sound, static starfield+male sound) by averaging responses across replicated trials.

For each treatment group per object size, we collected responses from a minimum of 45 subjects. In total, individual responses from 193 male and 95 female *An. coluzzi* has been used for this study.

### Free-flight data processing

We obtained a dataset from (*53*) comprised of six swarming events of male *An. coluzzii* mosquitoes, which provided us with 938 trajectories for analysis. The dataset captured the motion of each mosquito in a swarms of 13-27 individuals at a sampling rate of 50Hz in laboratory conditions, providing information on their three-dimensional locations over time. Additionally, we had at our disposal calculated velocity and acceleration vectors for each recorded frame. We analyzed this information using MATLAB R 2023a.

For each mosquito within the swarm, we identified its nearest neighbor at each frame and calculated flight parameters, such as acceleration towards nearest neighbor as well as magnitude of acceleration, speed, angular velocity, and flight direction relative to the position of the nearest neighbor. The acceleration towards nearest neighbor was determined by projecting the mosquito’s instantaneous acceleration onto the unit vector towards its nearest neighbor.

This measurement allowed us to infer the effective inter-individual forces, with positive values indicating attraction and negative values suggesting repulsion. Flight direction relative to the nearest neighbor was quantified by the angle between the mosquito’s velocity vector and the unit vector towards its nearest neighbor. Angular measures of 0°, 90° and 180° correspond to the mosquito flying directly towards, perpendicular to, and directly away from its neighbor, respectively. Lastly, we evaluated the angle between flight directions of nearest neighbors. This calculation sought to reveal the level of alignment in flight paths of interacting pairs of male mosquitoes as a function of nearest-neighbor distance.

To determine whether the observed flight dynamics were influenced by the presence of proximal neighbors or were a result of the general swarming motion, we implemented a time-shift analysis. In this analysis, each mosquito’s position at time *t* was virtually transplanted to the same position at time *t*+1 sec within the swarm. This transplantation allowed us to measure flight parameters of the same mosquito at *t* relative to its ‘virtual’ nearest neighbor at *t*+1sec and thus, compare the flight dynamics of mosquitoes in the presence of actual neighbors against those with the introduced ‘virtual neighbors’ in control scenarios. For this time-shifted analysis, we evaluated various time-shifted intervals beyond the initial 1-sec shift, and across different intervals, we consistently observed similar trends in the flight dynamics.

### Statistical Analysis

All statistical analyses were conducted using the R software, version 4.3.1. Given the large number of animals tested for each experiment, parametric tests were deemed suitable for statistical testing. Across all analyses, P-values for pairwise treatment comparisons were adjusted using the Holm method and a significant criterion of 0.05 was used for all statistical testing.

#### Response to rotating starfield patterns

To discern the influence of rotating starfield patterns on tethered mosquito behavior, we applied pairwise t-tests. These tests compared the subjects’ normalized steering responses (L-R WBA) and wingbeat kinematics (L+R WBA and WBFs) during the last second of rotating stimulus presentation against baseline responses recorded in the one-sec interval preceding the stimulus onset. We focused on the last second of stimulus presentation to account for the delay in responses. However, our results were consistent across various time intervals after accounting for the response delay.

#### Response to moving objects

When analyzing the visual response to a small moving object, we restricted our consideration to the initial 3 seconds of stimulus presentation, during which the object completed one turn across the visual field. This decision was informed by an observed perturbation point occurring at 3 seconds, where the object’s location reset, causing it to appear in the opposite visual field. This perturbation introduced variability in baseline responses, complicating the interpretation and analysis for the latter half of the stimulus duration. Responses to the full 6-sec stimulus are presented in the supplementary Materials.

To examine steering responses in these trials, paired t-tests were again used to evaluate changes in steering behavior by contrasting the last second of the stimulus cycle with the one second of pre-stimulus baseline. Although peak steering responses did not uniformly occur within the last second of stimulus cycle, results were consistent across various stimulus intervals after the initial 1.5 sec. Additionally, Spearman’s correlation analyses were performed to examine the relationship between the average responses and the object’s location during the 0.75 to 2.25-sec window, when the responses appeared to vary monotonically with object’s location.

Since aerodynamic thrust, measured through L+R WBA and WBF, appeared to modulate in relation to the object’s location, we performed two Spearman’s correlations assessing wing kinematics while the object was moving in front of the mosquito’s visual field (0-1.5 sec) and to when it was moving away (1.5-3 sec). Pairwise t-tests were then applied to determine whether the thrust responses during these distinct time frames, were statistically distinguishable.

Finally, to assess whether changes in flight responses compared to the baseline, denoted by Δ L-R WBA, ΔL+R WBA, and ΔWBF, were dependent on acoustic treatments and object sizes, we fitted linear mixed-effect models (LMMs) to account for repeated measurements. We fitted two nested LMMs for each of the three response variables. The first model included the fixed effects of acoustic treatments, object size as predictor variables and the random effect of subject ID. The second model included an additional interaction term of acoustic treatments × object size. ANOVAs were used to compare these nested models and thus, test if the interaction term was significant. The first model without the interaction term was adopted if no significant differences were found.

#### Free-flight analysis

To investigate the impact of proximity to a neighbor on flight dynamics, we stratified nearest-neighbor distances into discrete categories of 2 cm scale: distances under 2 cm formed one group, distances between 2 cm and 4 cm composed another group, and so forth. For each calculated flight parameter, we fitted a LMM model. This model included categorized distances, treatment conditions (real swarm versus the control with virtual neighbors) and their interaction as fixed effects. Given the longitudinal nature of our data, where flight parameters of each mosquito within a swarm were measured over time, we incorporated a nested random effect structure in our model. This structure accounted for repeated measures of individual mosquitoes within swarms, with time as a random slope to adjust for temporal variations in flight activity. Using this model, we compared the flight parameters between real swarm and control conditions across varying distances.

## Acknowledgements and Funding

We thank Simon Sawadogo for sharing pictures of natural swarms; Ruth Müller, Simon Sawadogo, Sofia Vielma, and members of the Riffell lab for feedback on this work; Binh Nguyen and Nicolas Avendano for helping with mosquito care; and Mridul Yadav for making the illustrations. This work was supported by grants from the Human Frontiers Science Program (HFSP -RGP0044/2021), National Institutes of Health (R01AI148300, R01AI175152), Air Force Office of Scientific Research (FA9550-21-1-0101, AWD-004055-G4) and French National Research Agency (ANR-15-CE35-0001-01)

