## Supplemental Information for "Multisensory integration in *Anopheles* mosquito swarms: The role of visual and acoustic information in mate tracking and collision avoidance"

Saumya Gupta *et al.*

**This PDF file includes:**

Figs. S1 to S6
Tables S1 to S5

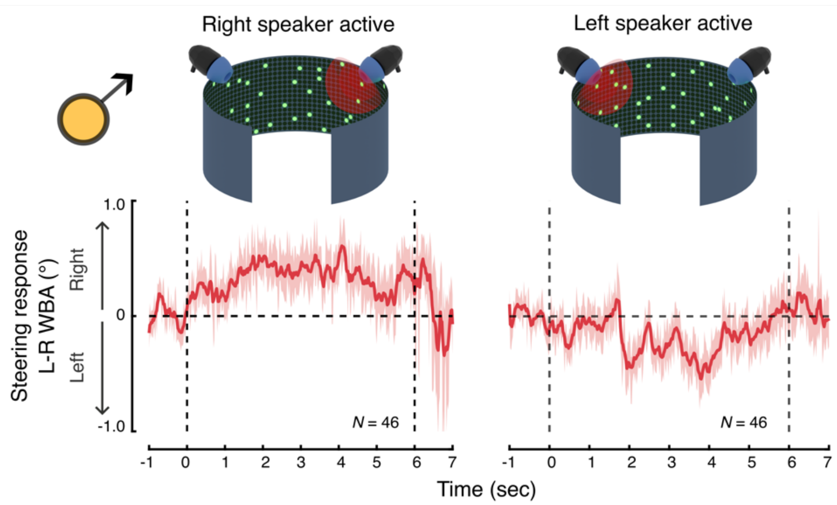

**Fig. S1. Phonotactic responses of males towards the location of female flight tones**. (Top row) Schematic representation of the audio setup within the flight simulator arena, illustrating the arrangement for two speakers positioned on the left and right side of the mosquito. (Bottom row) Mean normalized turning responses of males to female-like tone broadcast at 450 Hz from the right (column1) and left (column 2) sides relative to the tethered individual. Responses are presented for three intervals: 1 sec before, 6 sec during, and 1 sec after the tone is played. Shaded regions represent standard error (± SE). The data indicate that males consistently steer towards the active speaker—rightwards when the right speaker is active and leftwards when the left speaker is active. This acoustic-mediated steering response prompted us to position the speaker directly in front of the tethered mosquito for subsequent visual-acoustic experiments.

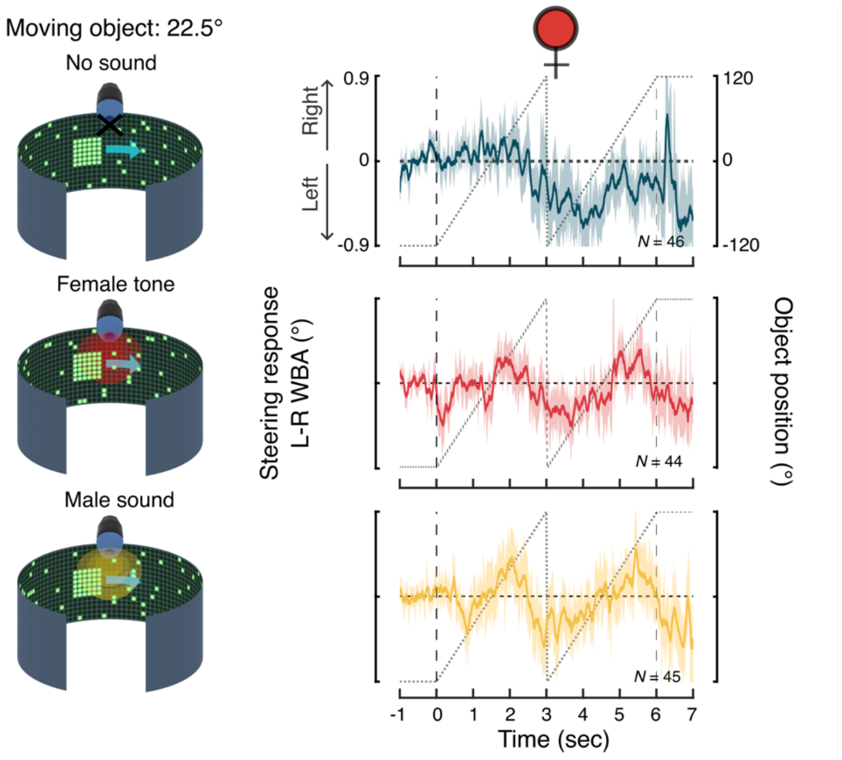

**Fig. S2. Extended analysis of female steering responses**. This figure illustrates the extended time series analysis of steering response by tethered female *An. coluzzii* (expanded from Figure 2). Here, the responses are shown for the full stimulus duration, capturing the mean behavior of the mosquitoes as they respond to a static object during the initial 1 sec (-1 to 0 sec), during two complete horizontal sweeps over the subsequent 6 sec (0 to 3sec and 3 to 6 sec), and in the final 1 sec (6-7 sec) when the object returns to a static position. The vertical demarcation at 3 sec indicates the perturbation point at which the object's position is reset.

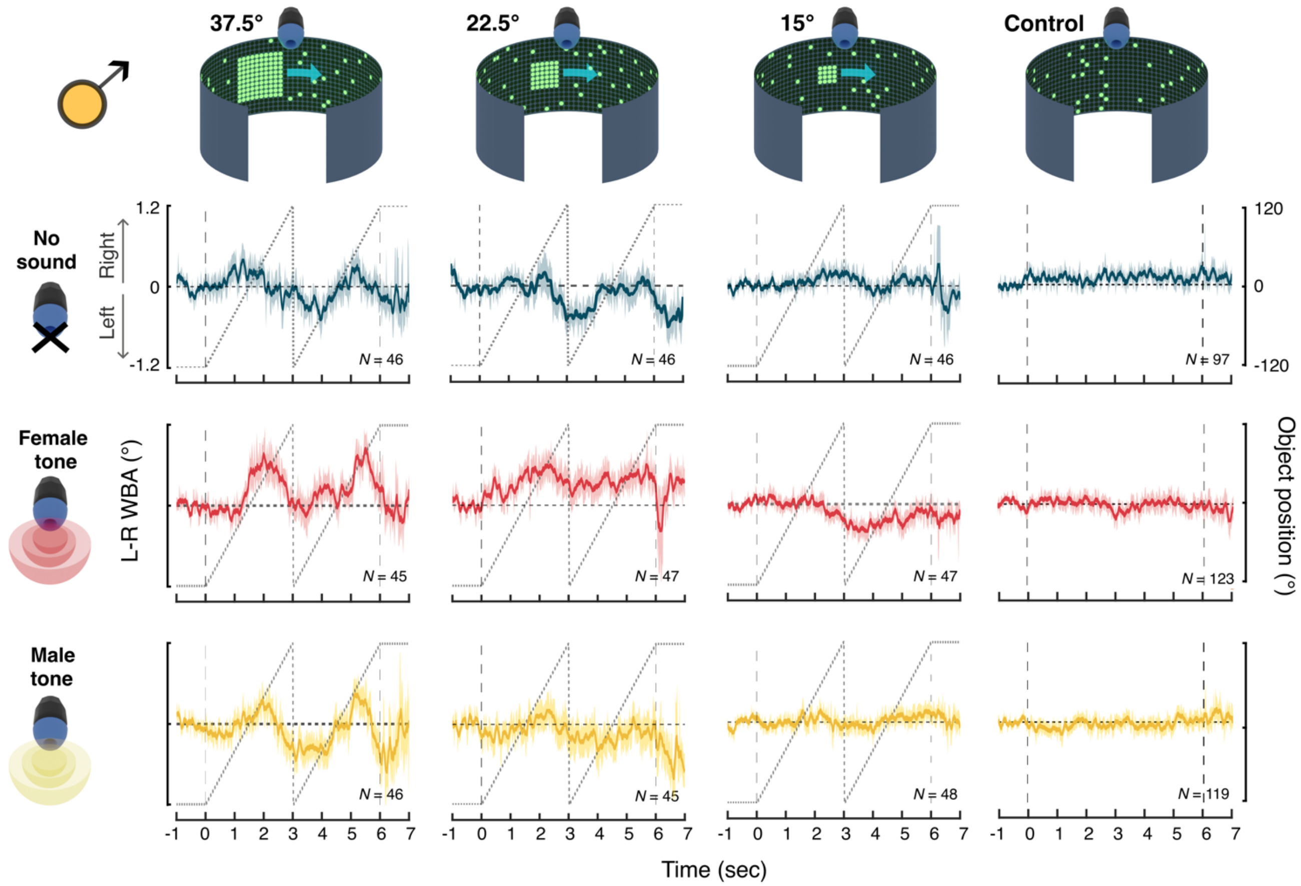

**Fig. S3. Male steering responses during extended stimulus presentation**. This figure illustrates the extended time series analysis of steering response by tethered male *An. coluzzii* (expanded from Figure 3). Here, the responses are shown for the full stimulus duration, capturing the mean behavior of the mosquitoes as they respond to a static object during the initial 1 sec (-1 to 0 sec), during two complete horizontal sweeps over the subsequent 6 sec (0 to 3sec and 3 to 6 sec), and in the final 1 sec (6-7 sec) when the object returns to a static position. The vertical demarcation at 3 sec indicates the perturbation point at which the object's position is reset.

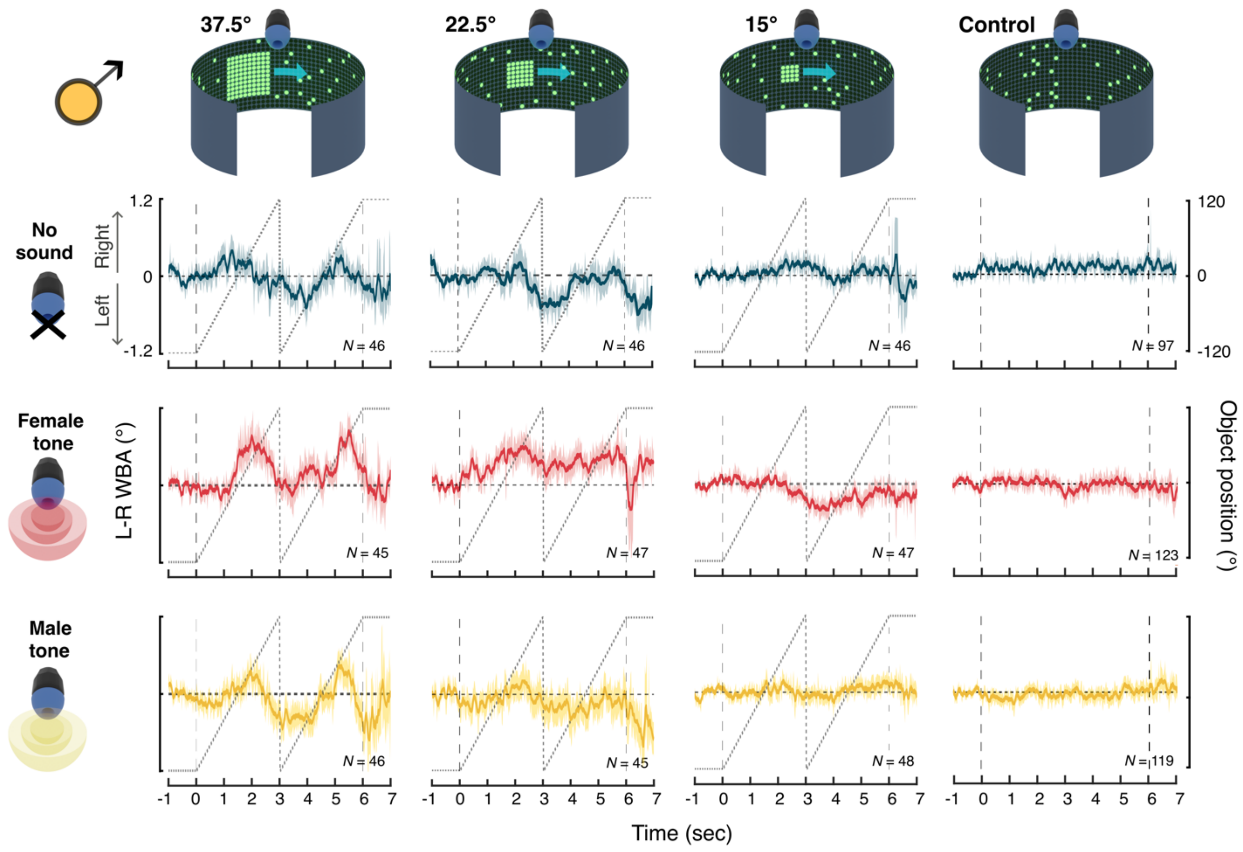

**Fig. S4. Male wingbeat amplitude during extended stimulus presentation**. This figure illustrates the extended time series analysis of the total wingbeat amplitude (L+R WBA) by tethered male *An. coluzzii* (expanded from Figure 4). Here, the responses are shown for the full stimulus duration, capturing the mean behavior of the mosquitoes as they respond to a static object during the initial 1 sec (-1 to 0 sec), during two complete horizontal sweeps over the subsequent 6 sec (0 to 3sec and 3 to 6 sec), and in the final 1 sec (6-7 sec) when the object returns to a static position. The vertical demarcation at 3 sec indicates the perturbation point at which the object's position is reset.

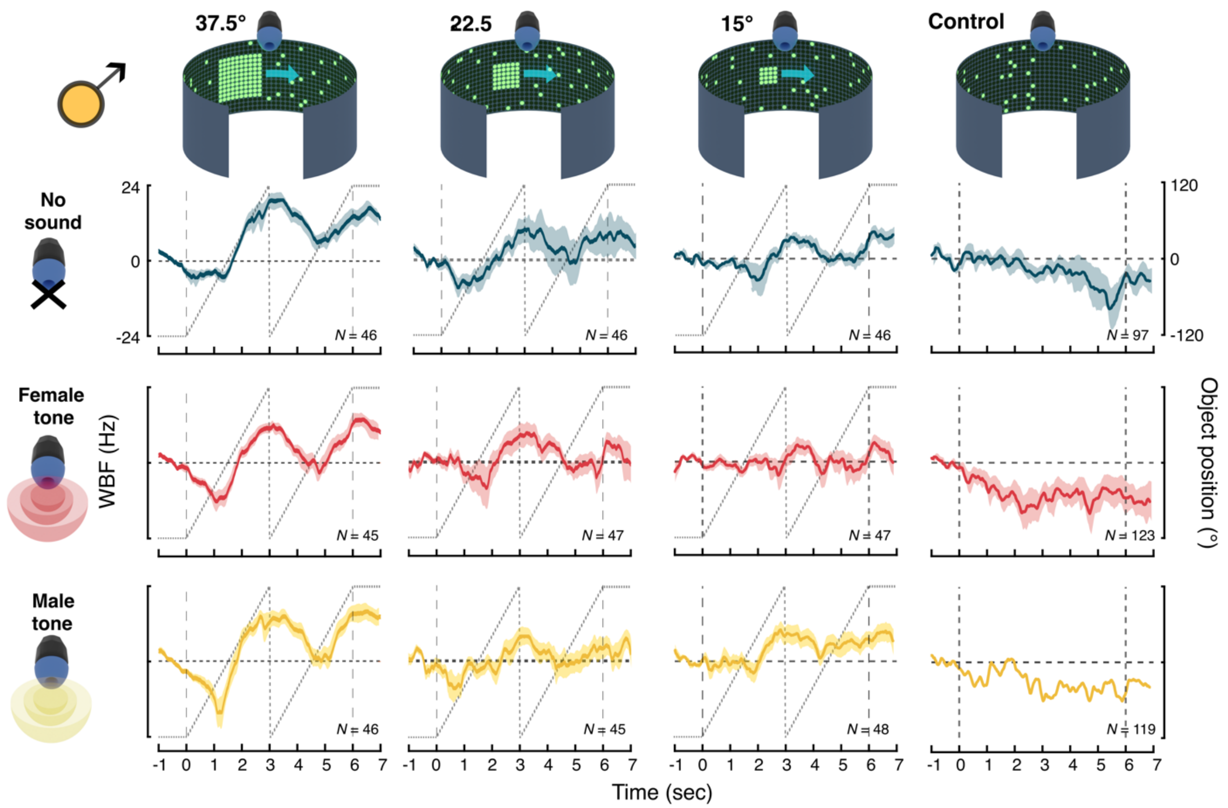

**Fig. S5. Male wingbeat frequency during extended stimulus presentation**. This figure illustrates the extended time series analysis of the wingbeat frequency (WBF) by tethered male *An. coluzzii* (expanded from Figure 4). Here, the responses are shown for the full stimulus duration, capturing the mean behavior of the mosquitoes as they respond to a static object during the initial 1 sec (-1 to 0 sec), during two complete horizontal sweeps over the subsequent 6 sec (0 to 3sec and 3 to 6 sec), and in the final 1 sec (6-7 sec) when the object returns to a static position. The vertical demarcation at 3 sec indicates the perturbation point at which the object's position is reset.

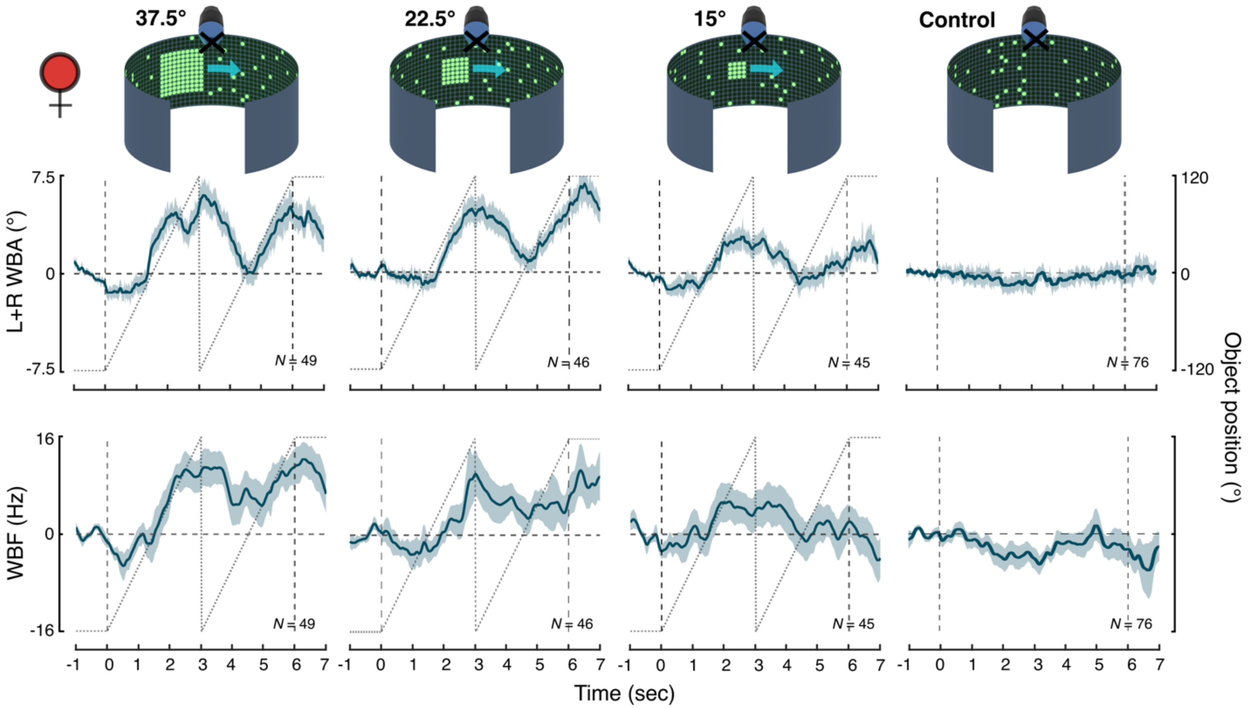

**Fig. S6. Female wing kinematics in response to objects of varying sizes**. This figure illustrates the mean normalized flight activity patters of females, as indicates by total wingbeat amplitude (L+R WBA; top panel) and wingbeat frequency (WBF; bottom panel) in response to objects of three different sizes and a control condition (static starfield). The responses are shown for the full stimulus duration, capturing the mean behavior of the mosquitoes as they respond to a static object during the initial 1 sec (-1 to 0 sec), during two complete horizontal sweeps over the subsequent 6 sec (0 to 3sec and 3 to 6 sec), and in the final 1 sec (6-7 sec) when the object returns to a static position. The vertical demarcation at 3 sec indicates the perturbation point at which the object's position is reset.

Table S1: Results of *linear mixed-effects model* (with significant interaction term) elucidating differential male steering responses to acoustic treatments (no sound, female tone, male tone) across visual conditions (37.5° object, 22.5° object, 15° object, and no object).

| | Estimate | SE | df | t ratio | *P* value |
|---|---|---|---|---|---|
| **37.5° object in static starfield background** | | | | | |
| *Female tone vs No sound* | *0.54* | *0.21* | *590* | *2.59* | *0.029** |
| *Female tone vs Male sound* | *0.47* | *0.21* | *590* | *2.28* | *0.046** |
| No sound vs Male sound | -0.06 | 0.21 | 590 | -0.32 | 0.750 |
| **22.5° object in static starfield background** | | | | | |
| *Female tone vs No sound* | *0.57* | *0.20* | *590* | *2.78* | *0.017** |
| Female tone vs Male sound | 0.41 | 0.21 | 590 | 2.01 | 0.091 |
| No sound vs Male sound | -0.16 | 0.21 | 590 | -0.76 | 0.450 |
| **15° object in static starfield background** | | | | | |
| Female tone vs No sound | -0.25 | 0.20 | 590 | -1.21 | 0.676 |
| Female tone vs Male sound | -0.11 | 0.20 | 590 | -0.55 | 1.000 |
| No sound vs Male sound | 0.14 | 0.20 | 590 | 0.67 | 1.000 |
| **Control: Static starfield background** | | | | | |
| Female tone vs No sound | -0.15 | 0.13 | 590 | -1.09 | 0.824 |
| Female tone vs Male sound | -0.02 | 0.13 | 590 | -0.12 | 0.901 |
| No sound vs Male sound | 0.13 | 0.14 | 590 | 0.97 | 0.824 |

Asterisks (*) denote significant differences ($P < 0.05$) between treatments when *P-values* are adjusted for multiple comparisons using the Holm method.

Table S2: Results of *linear mixed-effects model* (with significant interaction term) elucidating differential male steering responses to visual stimuli in the presence of female tone.

| **Treatment: Visual cues in the presence of female tone** | | | | | |
|---|---|---|---|---|---|
| | Estimate | SE | df | t ratio | *P* value |
| 37.5° vs 22.5° object | -0.04 | 0.21 | 590 | -0.21 | 1.000 |
| *37.5° vs 15° object* | *0.51* | *0.21* | *590* | *2.47* | *0.041** |
| *37.5° object vs Control* | *0.46* | *0.17* | *590* | *2.68* | *0.034** |
| *22.5° vs 15° object* | *0.55* | *0.20* | *590* | *2.71* | *0.034** |
| *22.5° object vs Control* | *0.50* | *0.17* | *590* | *2.98* | *0.018** |
| 15° object vs Control | -0.05 | 0.17 | 590 | -0.28 | 1.000 |

Asterisks (*) denote significant differences ($P < 0.05$) between treatments when *P-values* are adjusted for multiple comparisons using the Holm method.

Table S3: Results of *linear mixed-effects model* (without non-significant interaction term) elucidating the effect of visual stimuli and acoustic treatments on changes in male wingbeat amplitude (L+R WBA).

| | Estimate | SE | df | t ratio | *P* value |
|---|---|---|---|---|---|
| **Treatment** | | | | | |
| No sound vs Female tone | -0.08 | 0.25 | 596 | -0.32 | 0.899 |
| No sound vs Male tone | -0.27 | 0.25 | 596 | -1.06 | 0.876 |
| Female tone vs Male tone | -0.19 | 0.24 | 596 | -0.76 | 0.899 |
| **Visual Stimuli** | | | | | |
| *37.5° vs 22.5° object* | *1.44* | *0.35* | *596* | *4.10* | *< 0.001** |
| *37.5° vs 15° object* | *2.03* | *0.35* | *596* | *5.81* | *< 0.001** |
| *37.5° object vs Control* | *3.32* | *0.29* | *596* | *11.52* | *< 0.001** |
| *22.5° vs 15° object* | 0.59 | 0.35 | 596 | 1.69 | 0.092 |
| 22.5° object vs Control | 1.88 | 0.29 | 596 | 6.52 | < 0.001* |
| 15° object vs Control | 1.29 | 0.29 | 596 | 4.55 | < 0.001* |

Asterisks (*) denote significant differences (*P* < 0.05) between treatments when *P-values* are adjusted for multiple comparisons using the Holm method.

Table S4: Results of *linear mixed-effects model* (without non-significant interaction term) elucidating the effect of visual stimuli and acoustic treatments on changes in male wingbeat frequency (WBF).

| | Estimate | SE | df | t ratio | *P* value |
|---|---|---|---|---|---|
| **Treatment** | | | | | |
| No sound vs Female tone | 0.79 | 1.74 | 596 | 0.455 | 1.000 |
| No sound vs Male tone | 1.12 | 1.75 | 596 | 0.640 | 1.000 |
| Female tone vs Male tone | 0.33 | 1.69 | 596 | 0.192 | 1.000 |
| **Visual Stimuli** | | | | | |
| *37.5° vs 22.5° object* | *11.10* | *2.47* | *596* | *4.48* | *< 0.001** |
| *37.5° vs 15° object* | *17.46* | *2.46* | *596* | *7.09* | *< 0.001** |
| *37.5° object vs Control* | *22.24* | *2.02* | *596* | *11.03* | *< 0.001** |
| *22.5° vs 15° object* | *6.37* | *2.46* | *596* | *2.59* | *0.020** |
| *22.5° object vs Control* | *11.14* | *2.02* | *596* | *5.51* | *< 0.001** |
| *15° object vs Control* | *4.77* | *1.99* | *596* | *2.40* | *0.020** |

Asterisks (*) denote significant differences (*P* < 0.05) between treatments when *P-values* are adjusted for multiple comparisons using the Holm method.

Table S5: Results of *linear mixed-effects model* (with interaction term) elucidating the differences between flight parameters of mosquitoes in real swarm versus control scenario at different nearest neighbor distances.

| | Distance | Estimate | SE | df | t ratio | *P* value |
|---|---|---|---|---|---|---|
| **Acceleration** | | | | | | |
| | *0-2 cm* | *0.23* | *0.04* | *615346* | *5.25* | *< 0.001** |
| | *2-4 cm* | *0.09* | *0.02* | *615346* | *5.10* | *< 0.001** |
| | 4-6 cm | 0.01 | 0.01 | 615346 | 0.96 | 0.339 |
| **Angular velocity** | | | | | | |
| | *0-2 cm* | *38.37* | *5.68* | *613071* | *6.75* | *< 0.001** |
| | *2-4 cm* | *9.57* | *2.23* | *613071* | *4.28* | *< 0.001** |
| | 4-6 cm | 0.070 | 1.47 | 613071 | 0.47 | 0.636 |
| **Flight speed** | | | | | | |
| | 0-2 cm | -0.003 | 0.002 | 615346 | -1.16 | 0.247 |
| | 2-4 cm | 0.001 | 0.001 | 615346 | 0.92 | 0.358 |
| | 4-6 cm | 0.0002 | 0.0007 | 615346 | 0.26 | 0.791 |
| **Flight direction relative to its nearest neighbor** | | | | | | |
| | 0-2 cm | -0.64 | 0.82 | 615346 | -0.08 | 0.431 |
| | 2-4 cm | -0.31 | 0.32 | 615346 | -0.97 | 0.331 |
| | 4-6 cm | -0.28 | 0.21 | 615346 | -1.32 | 0.186 |
| **Angle between flight directions of neighboring mosquitoes** | | | | | | |
| | *0-2 cm* | *-5.89* | *1.06* | *350950* | *-5.54* | *< 0.001** |
| | *2-4 cm* | *-0.94* | *0.42* | *350950* | *-2.25* | *0.024** |
| | *4-6 cm* | *-0.81* | *0.28* | *350950* | *-2.928* | *0.003** |

Asterisks (*) denote significant differences ($P < 0.05$) between data from real swarm and control.
